# Low-Cost 3D-Printed Tools Towards Robust Longitudinal Multi-Modal Pre-Clinical Imaging

**DOI:** 10.1101/2023.12.07.570719

**Authors:** Nader Allam, Edward Taylor, I. Alex Vitkin

## Abstract

Window chamber models enable a range of preclinical *in-vivo* optical studies with high spatial resolution and contrast, most notably probing the tumour microenvironment (TME). However, there are multiple sources of experimental variability that can affect the quality of the resultant data, especially in the context of longitudinal data acquisition, where accurate registration between images acquired at different times is crucial to understanding changes to the spatially heterogeneous TME. Further, it is challenging to correlate the findings of these models to clinical imaging modalities such as magnetic resonance imaging (MRI), which typically have much lower resolution and derive contrast from different physical mechanisms; yet such correlations may assist in the translation of window-chamber-derived basic preclinical insights into the clinic. Here, we describe the development and construction of a low-cost 3D-printable window chambers and compatible toolset to improve the accuracy, precision, and repeatability of longitudinal pre-clinical imaging and inter-modality co-registration at different spatial resolution scales. Such improvements in our novel multi-modal experimental pipeline may assist researchers in the acquisition and translation of TME biomarkers and other pre-clinical measurements from the window chamber model into the clinic.

## 2 Introduction

First developed in the 1930s, surgically implantable window chambers (WCs) enable high-resolution optical interrogation of tissue microstructure and its vasculature, with wide-ranging applications across the fields of biophysics, biomaterials, and biomedical engineering and biomedicine^1–16^. In particular, WC models have been utilized to shed light on the tumour microenvironment (TME), such as the microvasculature which is hypothesized to play a major role in pathology development and tumour therapeutic response^4–10^. They consist of a surgical opening into the tissues of an organism (most commonly small rodents such as mice or rats), sealed by a biocompatible frame (typically made of titanium) securing a glass window over the target organ to be imaged^17^. In rodent WC models, the loose skin in the back (dorsal side of the animal) is often selected for WC installation due to its convenience for imaging with reduced motion artifacts; that said, cranial, spinal, abdominal and other WC preparations, although less common, are also possible^14–16^. But the *dorsal skinfold WC* (DSWC), in which the inner sides of the skinfold (the panniculus carnosus muscle, the final layer underneath the subcutis) is exposed for imaging^1^ (see **Figure 1**), remains the most widely used WC in biomedical research.

**Figure 1:**
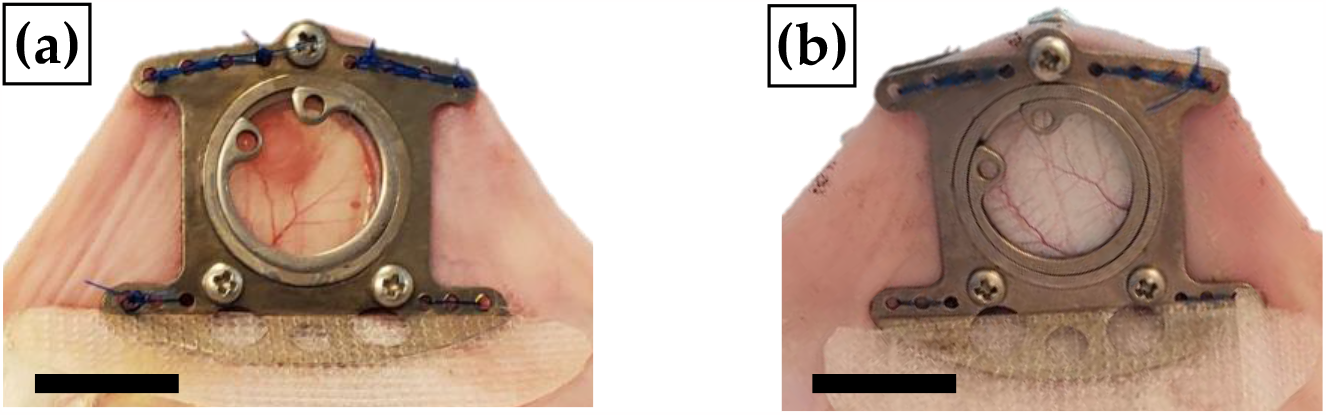
Typical titanium dorsal skinfold window chamber (DSWC) model. These DSWCs were manufactured by APJ Trading Co. (Ventura, CA, USA) and surgically installed on (a) tumour bearing (b) bare-skin (healthy) NRG mice. Here a ∼100 μm gap separates the glass window from the exposed panniculus carnosus muscle layer. Scale bar = 1.0 cm.

Although the basic DWSC design is now quite mature, improvements can be made to optimize quantification of therapeutic response assessment and translatability of DWSC optical signals to widely used oncological imaging modalities such as magnetic resonance imaging (MRI) and computed-tomography (CT). DWSC designs should be optimized to ensure both inter-timepoint and inter-modality imaging spatial co-registration, as well as WC mouse longevity for the necessary long(er)-term post-treatment response assessment studies (on the order of weeks-months).

While the DSWC model was designed to facilitate longitudinal optical imaging experiments in a preclinical setting, the translation path towards the clinic is limited. One possible route is to correlate the details of the WC findings with the coarser images and metrics obtained with imaging modalities that *are* used in the clinic, such as MRI and CT. This way, these can be infused with additional detailed insights from the longitudinal optical investigations, thus enabling a form of clinical translation for the WC studies. For example, due to its superior soft-tissue contrast, MRI is widely used in oncology to assess and stage the pre-radiotherapy (RT) tumours, as well as for post-RT response monitoring^18–20^. CT is also employed in radiation oncology for treatment planning, patient set-up verification, and occasionally treatment response assessment ^21–24^. However, the significant magnetic susceptibility and high atomic number of the titanium metal (relative to tissue) used in many DSWC frames will inhibit any optical-to-MRI/CT correlative studies. Some researchers have begun experimenting with various polymers including 3D-printed plastics and silicone for the WC construction to enable longitudinal multi-modal imaging studies, however with arguably low accuracy and precision of inter-modality spatial co-registration due to the lack of any clearly-defined fixed fiducial markers visible across the employed imaging modalities^11,25^. Motion artifacts in all imaging modalities (primarily respiration-related) and lack of temporal repeatability in data acquisition over the timeframe of several weeks create challenges for longitudinal tracking of regions of interest. DSWC design improvements addressing these problems may thus increase image quality and robustness of findings, and perhaps facilitate the clinical translatability of intravital optical measurements.

The DSWC installation presents another opportunity for improvement. Specifically, reducing the chances of infections and tissue necrosis (causing transudate and exudate accumulation) is important for maintaining optical clarity, and reducing the risk of DSWC tilt resulting from the surgical implant procedure may improve mouse longevity and thus potential experiment duration. Certainly obtaining better data for longer times would be highly valuable for DSWC research, in the context of RT response quantification and beyond.

Here we present a comprehensive experimental pipeline to address these various identified shortcomings. Our proposed solution describes the development of a fully resin-printed DSWC and a peripheral toolset, both addressing the sources of variability towards improving accuracy, precision and repeatability of pre-clinical longitudinal multi-modal imaging experiments.

## 3 Methodology

### 3.1 Animal care and ethics

All animal handling procedures were approved by the University Health Network’s institutional Animal Care Committee in accordance with the guidelines of the Canadian Council for Animal Care (AUP #3256).

Proper measures were taken to minimize animal discomfort through administration of anesthesia and analgesia as necessary in doses recommended by veterinarians of the University Health Network’s Animal Resources Centre. Analgesia consisted of SR buprenorphine, 1 mg/kg, 0.6 mg/mL in 0.9% saline immediately prior to surgery, and Metacam, 5 mg/kg, 0.5 mg/mL, immediately post-surgery and every ∼24h for 48-72 hours. Anesthesia consisted of 5% isoflurane with 1 L/min O_2_ flowrate for induction and ∼1-2% isoflurane (as a function of mouse state of consciousness) with 0.5 L/min O_2_ for maintenance for all procedures except for sacrifice whereas a cocktail mix of 100 mg/kg, 10 mg/mL ketamine, and 10 mg/kg, 1 mg/mL xylazine in 0.9% saline was administered to induce anesthesia in normoxic conditions.

### 3.2 Design of DSWC and peripheral toolset

The target design parameters were chosen based on a literature review of material compatibility with various imaging modalities and the most commonly reported sources of biological and experimental variability in the DSWC model, with further empirical refinements upon over the multiple design iterations (see **<u>Suppl. 1</u>**). The requirements the DSWC and associated toolset must fulfill are simplified as follows:

i. <u>Material safety</u>: The DSWC should be biocompatible, sterilizable^26^, and “MR-safe” (non-ferromagnetic and with no metallic components in contact with the mouse to avoid heating injuries).
ii. <u>Maintenance of optical clarity</u>: Weeks following window chamber installation, transudate (∼transparent/translucid) and exudate (highly turbid) are released, crucially obscuring the visualization afforded by optical microscopies^25,27,28^ This may be a result of elevated cytokine activity as an inflammatory response to the skin wound, noted in all WC installations^29^. It has been demonstrated that if an air gap is present over the exposed panniculus carnosus muscle tissue, exudate formation is exacerbated.
iii. <u>Ease of imaging and inter-timepoint co-registration</u>: Towards robust longitudinal imaging, it is important that motion artifacts be kept minimal and that timepoint to timepoint imaged field-of-view (FOV) be kept consistent, given the overarching objective of gathering quality data for time series analysis.
iv. <u>Ease of inter-modality correlations</u>: The frame should have design features facilitating inter-modality co-registration (e.g., fiducial markers visualizable with relevant imaging modalities, etc), both for optical-MRI and optical-histology correlations.

All parts were prepared in Fusion 360^®^ computer-aided-design (CAD) software (Autodesk Inc.).

### 3.3 Fabrication of DSWC and peripheral toolset

To satisfy criterion I., the DSWC was designed and fabricated with properties exhibiting high temperature-resistance for autoclaving / sterilization, low atomic number and irradiation compatibility (specifically for CT imaging and RT response monitoring studies, respectively), magnetic susceptibility similar to tissue and finally effective removal of the air gap from the DSWC installation procedure. Thus, we chose to 3D-print the WCs using fusion deposition modeling (FDM) and stereolithography apparatus (SLA), leveraging the material properties of polymer plastics^30–33^. All DSWC parts except for the hula-clip (defined in **sec. <u>4</u>**.**<u>1</u>**) were printed out of Formlabs BioMed Clear Resin^®^ (Formlabs inc.), a USP Class VI certified material designed for long-term skin and mucosal membrane contact, with heat resistance up to ∼130°C, printed with 50 μm layer-height ^34^ via the Elegoo Mars 3 masked SLA (MSLA) printer. To maximize optical and near-infrared wavelength range transparency, the front frame SLA printed part was sprayed with USC SprayMax 2K Glamour High Gloss Aerosol Clear^® 35–38^. As this spray coat is not designed to be biocompatible, the backside of the front-frame was masked by an off-the-shelf polyvinyl chloride tape (PVC “packing tape”) during the spray-coat process.

The hula-clip does not require as high resolution printing but needs very high durability at a low weight; it was therefore FDM printed on a Prusa^®^ MK3S out of a high strength and durability blend consisting of nylon copolymer, reinforced with carbon-fiber (CF) (heavy-duty CF-nylon filament^®^, Filaments Inc, Toronto, ON, CA), with heat resistance up to ∼167°C. The full DSWC weighed up to ∼0.9 g, less than 50% of most other commercially available DSWCs; this is advantageous for longitudinal imaging ^11,12^ over the weeks/months time frame as desirable in treatment response monitoring studies.

### 3.4 DSWC surgical installation

A surgical stage and “surgical guide” were designed to enable this full DSWC installation procedure to be performed under isoflurane anesthesia and with improved control to maximize precision. The stage consists of a custom-designed heated mouse-bed with isoflurane inlet tubes fitted onto a third-arm soldering stage (Kotto^®^)^39^ meant to help manipulate and fix the “surgical guide” for accurate positioning of the skin during DSWC installation. In other words, it facilitates the determination of the ideal position for subcutaneous tumour inoculation on the mouse flank and for DSWC installation, 3-4 weeks later. This ensures that the skinfold can be fixed comfortably upright on the mouse’s back^11^ while the tumour grows centered within the FOV of the DSWC (see **Figure 2**). Please refer to **<u>Suppl. 2</u>** for full details on the surgical protocol and **<u>Suppl. 3</u>** for details of fabrication of the surgical stage.

**Figure 2:**
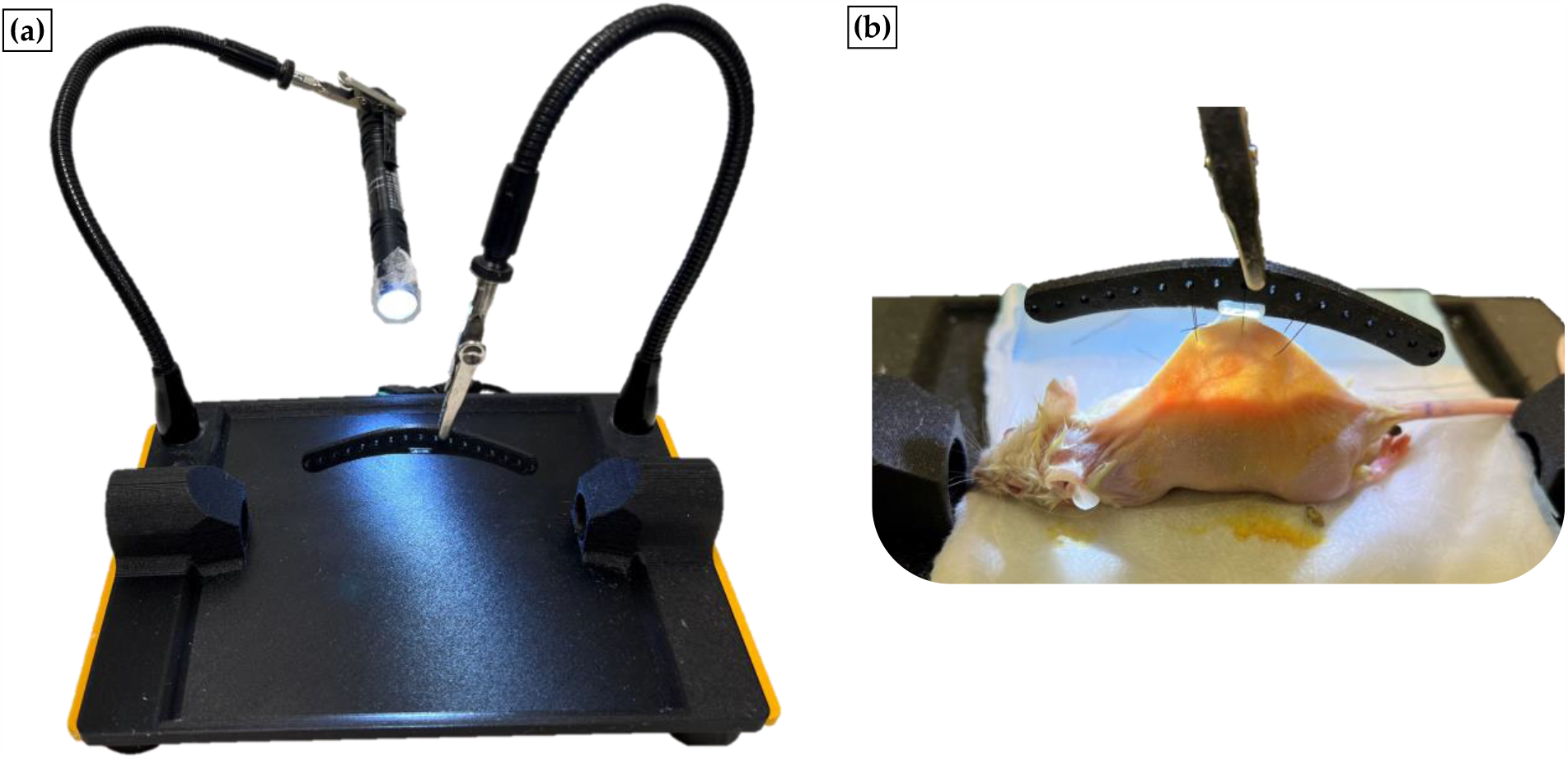
Isoflurane enabled DSWC installation. (a) represents the surgical stage on its own and (b) in practice, assisting the surgeon in holding the back DSWC frame via the “surgical guide” tool as well as a flashlight to transilluminate the skinfold and further guide accurate positioning of DSWC front frame. Please refer to **<u>Suppl. 2</u>** for the detailed surgical protocol. All figures were prepared in Fusion 360.

### 3.5 Imaging

All experimental data acquisition began ∼5 days post-DSWC installation to allow mice to recover from the surgery (while some studies report this inflammatory response period can last as little as a day^11^, several others suggest that it may take up to 3-5 days^27,40^), minimizing the impact of confounding variables pertaining to tissue inflammation such as the secretion of the largely anti-angiogenic Angiopoietin-2 ^40,41^ . These would otherwise impact the window chamber optical clarity and the natural state of the imaged vascular architecture *In-vivo* imaging was performed optically (via brightfield microscopy and optical coherence tomography (OCT)), via MR, via CT and finally *ex-vivo* via histology. Full details regarding optical and MR imaging protocol may be found in our previous publication^8^.

## 4 Results & Discussion

### 4.1 Description of DSWC

With criterion I. addressed via the material choice as described in section **3.3**. The final design of the DSWC consisted of (refer to **Figure 3**Error! Reference source not found.(d)):

- A combined window (11 mm diameter, 100 *μm* thick) and circular front frame (17 mm diameter, 1.7 mm thick with fully filleted outer rim), printed as a single part. This was designed to be installed underneath the skin, similar to the installation protocol of many orthotopic WC^25^, in an attempt to reduce chances of an air gap between the tissue and the window, to mitigate sources of infection and improve tissue fixation in the DSWC, thus addressing criterion II (see sec. **3.2**). Reduction of the air gap reduces the chance of magnetic susceptibility artifacts in MRI^42^. The front frame also includes 4 extrusions enabling interfacing with various tools facilitating longitudinal multi-modal imaging.
- Back frame (11 mm wide and 21 mm tall) with 3 spacers (installed separately), a bottom 45° flange to help keep the DSWC dorsal skinfold upright (important for preventing changes in tissue perfusion longitudinally as well as mouse comfort^11,12^) and a top extrusion for interfacing with imaging and treatment stages.
- Hula-clip support (35 mm long and 10 mm wide) resting on the mouse’s back (thus applying minimal torque to the skinfold) enabling a more compact and lighter DSWC design all the while helping to keep the skinfold upright, by fitting into the back frame and wrapping around the skinfold toward the front frame.
- Chamber protector (19 mm wide and 14 mm tall) pressure fitted onto the front frame extrusions acting to protect the window from dirt accumulation and damage from the mouse daily activities as well as the sutures securing the front frame to the back frame from the mouse gnawing. This cover is meant to be replaceable, only removed during imaging and treatment procedures whereas alternative tools may be added as later described.

**Figure 3:**
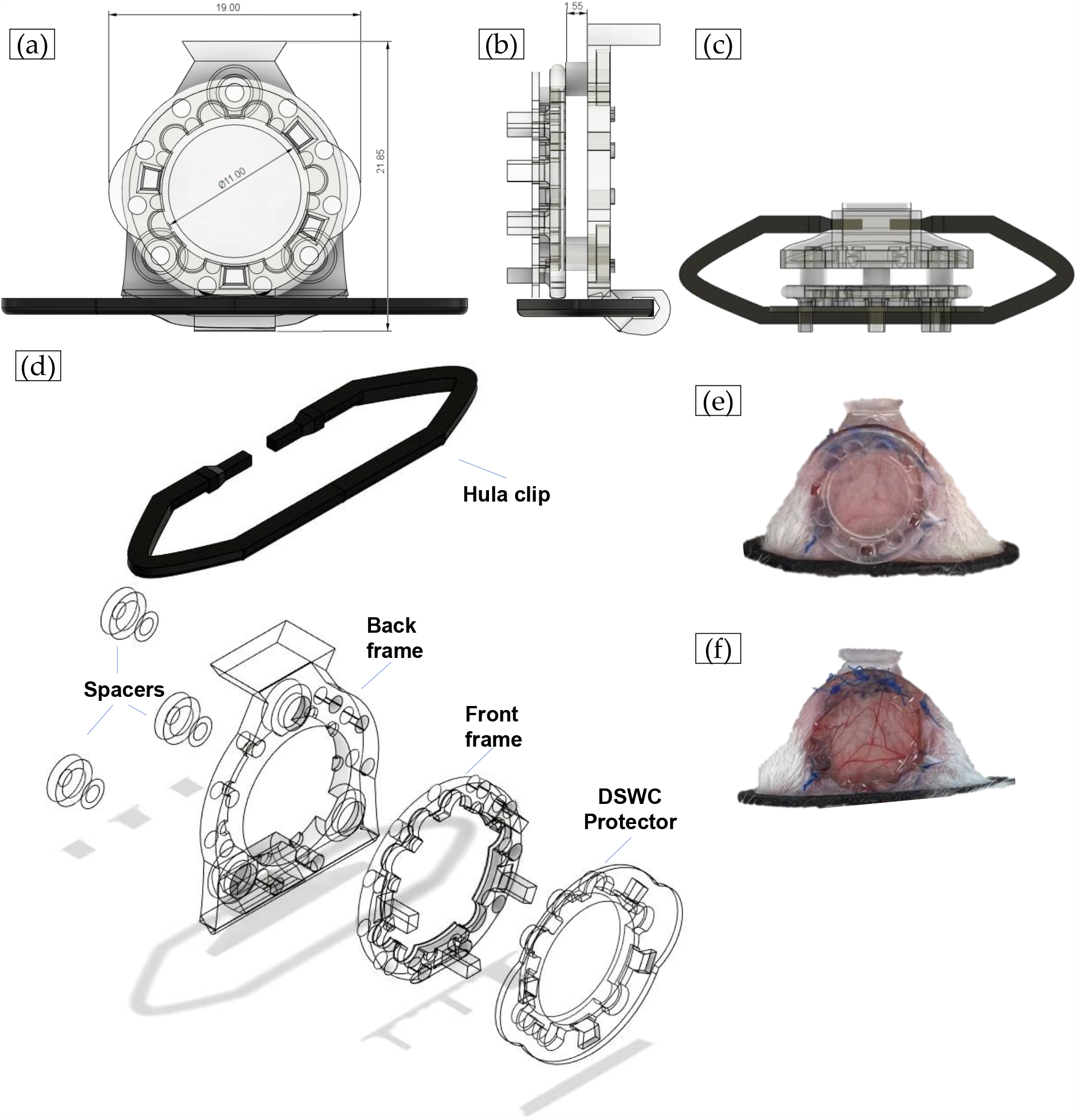
Novel fully resin 3D-printed DSWC. (a) Coronal, (b) Sagittal, and (c) Top transverse views of the design. (d) represents the individual components of the DSWC to be assembled. Finally a zoomed-in view of the DSWC in practice is represented on an anesthetized NRG mouse (e) with and (f) without the DSWC protector secured. All figures were prepared in Fusion 360.

### 4.2 Description of Peripheral toolset: Longitudinal and multi-modal imaging

To address criteria II. and III., representing the design features enabling robust longitudinal imaging co-registration and robust multi-modal imaging co-registration respectively, various imaging (and treatment) beds/restrainers and alternative DSWC frame peripheries were designed, designated as the “peripheral toolset”. The imaging and treatment beds/restrainers serve dual functions: motion artifact mitigation and improvement of inter-timepoint DSWC positioning reproducibility. The first is self-explanatory for its well-known degradative effects on imaging quality^43–46^. The latter is crucial for 2 reasons: firstly, the phenomenon of exudate accumulation (described in DSWC criteria I. above); and secondly, facilitating sub-region-specific longitudinal monitoring important for monitoring metrics sensitive to the microvascular heterogeneity^8,9^. While techniques for addressing both these issues computationally have been developed^9,47–49^, it is usually ideal that such issues be addressed prior to the data acquisition given the relatively lower cost and higher potential effectiveness of such hardware solutions. The peripheral toolset act to facilitate inter-modality spatial co-registration across different resolution scales between optical intravital microscopy and MRI as well as between the former and histological sections. These solutions detailed below are represented in **Figure 4** and **<u>Suppl. 3</u>**:

**Figure 4:**
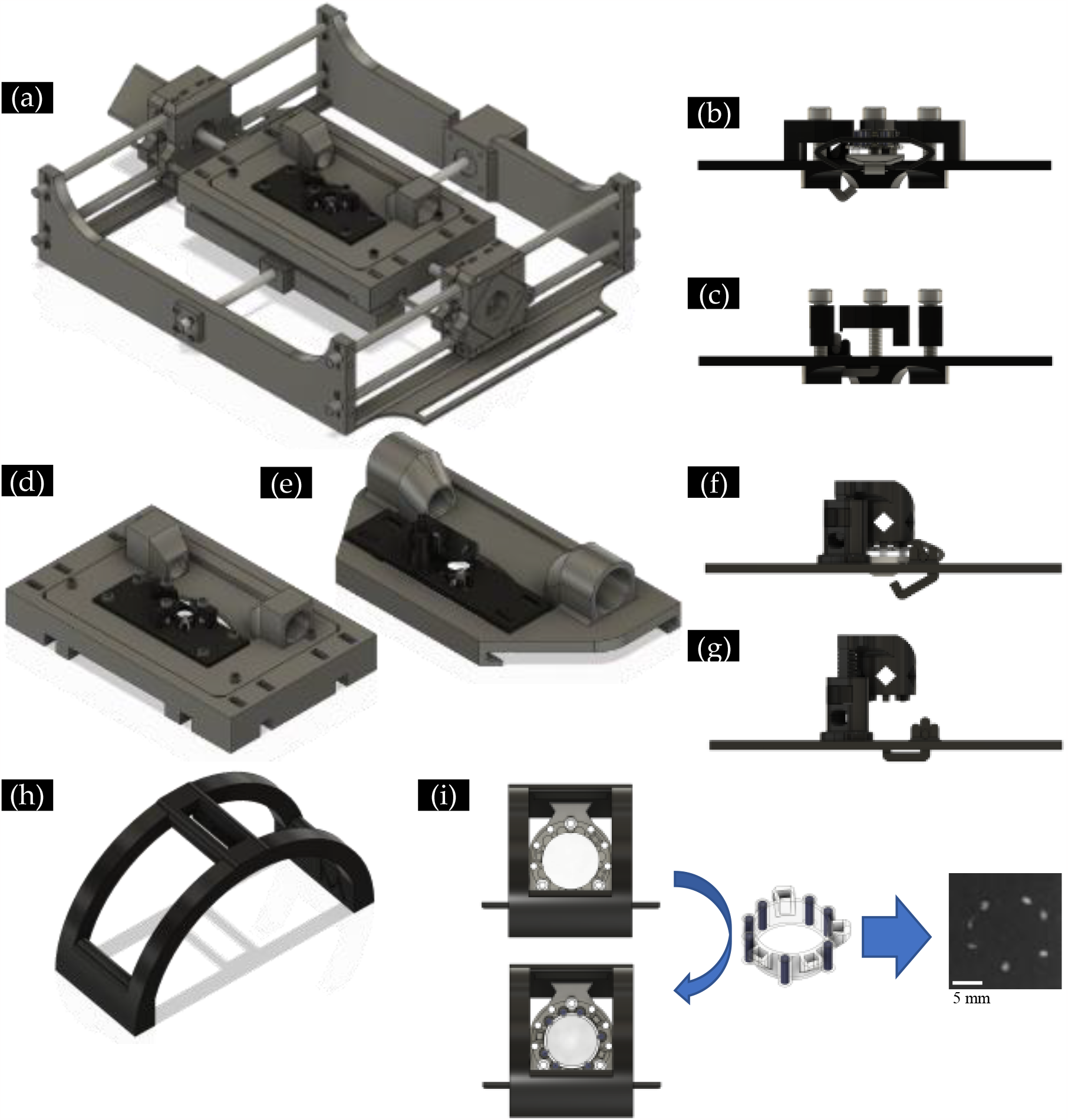
Custom-designed peripheral toolset enabling more robust longitudinal and multimodal DSWC imaging. (a) Automatic xy-microstage (excluding electronics) with metal optical imaging & treatment (MOIT) restrainer. (b) MOITR in configuration securing a DSWC.(c) MOITR in its loosened configuration. (d) Portable stage for use in setups non-conducive to the automatic xy-microstage.(e) CT clampable stage with all-plastic optical imaging & treatment restrainer (POITR) for use in rotating gantry setup, such as an irradiator or CT scanner. (f) POITR in configuration securing a DSWC. (g), POITR in its loosened configuration. (h) MRI-bore restrainer for use in MRI scanner. (i) MRI-bore restrainer keeping the DSWC static in an upright orientation, while the fiducial markers ring is slipped onto the front frame. Please refer to **<u>Suppl. 5</u>** for a video file showcasing the window chamber design and its interfacing with the various peripheral tools. Scale bar represents 5 mm. All figures were prepared in Fusion 360.

For robust longitudinal optical imaging a custom-designed, 3D-printed, assembled and programmed xy-microstage mouse restrainer was made^50^. The frame was printed out of a combination of elastomer plastic (thermoplastic polyurethane) and a rigid plastic, polylactic acid to strengthen the stage all the while dampening any vibrations from the motors over the course of imaging. Mice are laid-down anesthetized via isoflurane on a heated bed (via nichrome wire array) with window chambers fixed through a custom-designed restrainer piece for the DSWC with adjustable tension toward avoiding oversaturation of backscattered light intensity through incorrect tilting of the chamber underneath the imaging probe. A pair of Nema17 stepper motors with 1.8°/step phase control connected to a threaded rod of 8 mm diameter and 4 mm pitch were each controlled by separate A4988 stepper motor controllers receiving commands from a Keeyee’s Arduino Uno^®^ clone microcontroller. The system also contains mechanically actuated end-stops as part of calibration positioning sequence (“homing”). This setup allows for positioning precision of 2.5 *μm* and using the GRBL interface “Universal G-code Sender”, the user may input g-codes to reproduce imaging positions inter-timepoint with high fidelity. These automatic micro-stages remove the problem of having to figure out the mouse positioning for each imaging timepoint, saving valuable time and removing imaging variability in these logistically complex experiments (see **Figure 5**). This also facilitates to first-order the inter-timepoint co-registration required for time series analyses^5,7,9^ and inter-modality co-registration from intravital optical imaging to ex-vivo histology (see **<u>Suppl. 4</u>**). Please refer to **<u>Suppl. 6</u>** for a video file representing the application of the xy-microstage.

**Figure 5:**
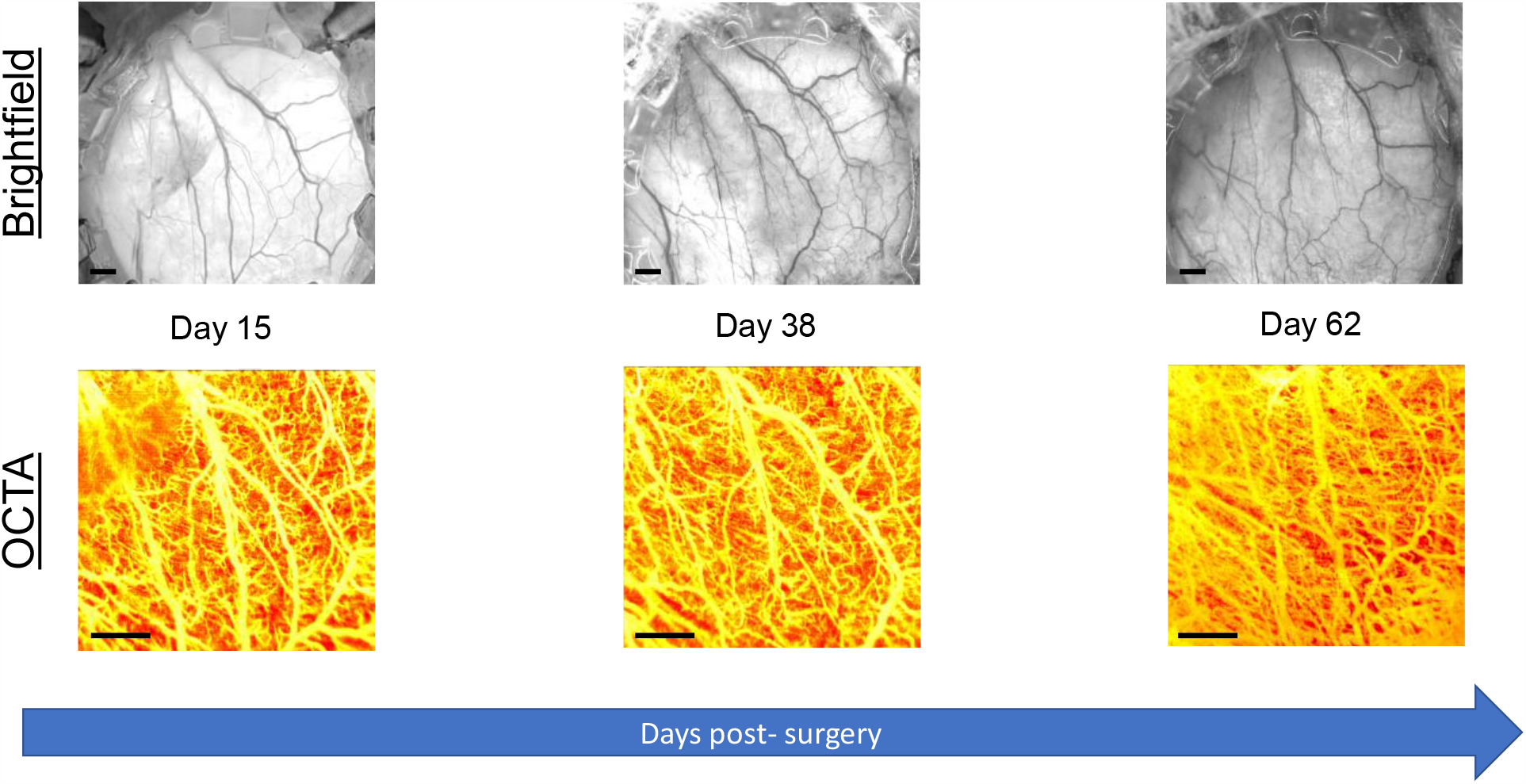
Longitudinal imaging of the novel DSWC installed on a bare skin mouse. The imaging is performed up to approximately 2-2.5 months post-installation of DSWC via various modalities such as brightfield and OCTA using the restrainers described in this section and shown in **Figure 4**. The scale bars represent 1 mm.

Seeking to create a truly multi-modal imaging pipeline, alternative mobile versions of the optical imaging restrainer bed were designed to restrain motion and improve positioning repeatability in a range of spatial constraints. These range from transillumination brightfield and epifluorescence imaging (**Figure 4**(d)) X-ray CT imaging (**Figure 4**(e)), and MR imaging (**Figure 4**(h)).

Given the increasing clinical relevance of MR imaging due to superior soft-tissue contrast over CT imaging, particular interest lies in bridging the resolution gap between MRI and high resolution optical imaging with limited penetration depth such as OCT to identify correlations and thus potentially improve clinical translatability of metrics extracted from the latter. for the monitoring of vascular perfusion further facilitate MR imaging custom-designed 3D printed CF-Nylon blend window chamber immobilization device reduced respiratory movement and aligned the window chamber along the sagittal plane in the MRI bore. A replaceable “fiducial marker ring” was placed on the mouse front frame prior to imaging to facilitate later MR image co-registration to OCT. It consists of 7 ∼0.8 mm diameter and ∼1.6 mm deep cylindrical wells filled with a mixture of tear-gel and a few drops of blue food-dye, tightly sealed by a drop of resin.

These wells thus act as both proton-dense fiducials appearing brightly in T2-weighted imaging^51^ and high contrast fiducials for the brightfield imaging. This tool is designed to interface with the front frame extrusions (see **Figure 3** and **<u>Suppl. 5</u>**) thus removing uncertainty in its positioning inter-modality (see **Figure 4**). The pipeline for inter-modality co-registration is visually described in **Figure 6** from MRI (and CT) to OCT. MRI was first co-registered to the brightfield image based on the fiducial marker ring, which was in turn co-registered to the OCT acquisition (all laterally) through a series of geometric similarity transformations, guided via manually identified fiducials with MATLAB built-in functions. During MR data acquisition, captured slices were specially chosen to align with the tissue surface, thus aligning axially with OCT-defined VOI.

**Figure 6:**
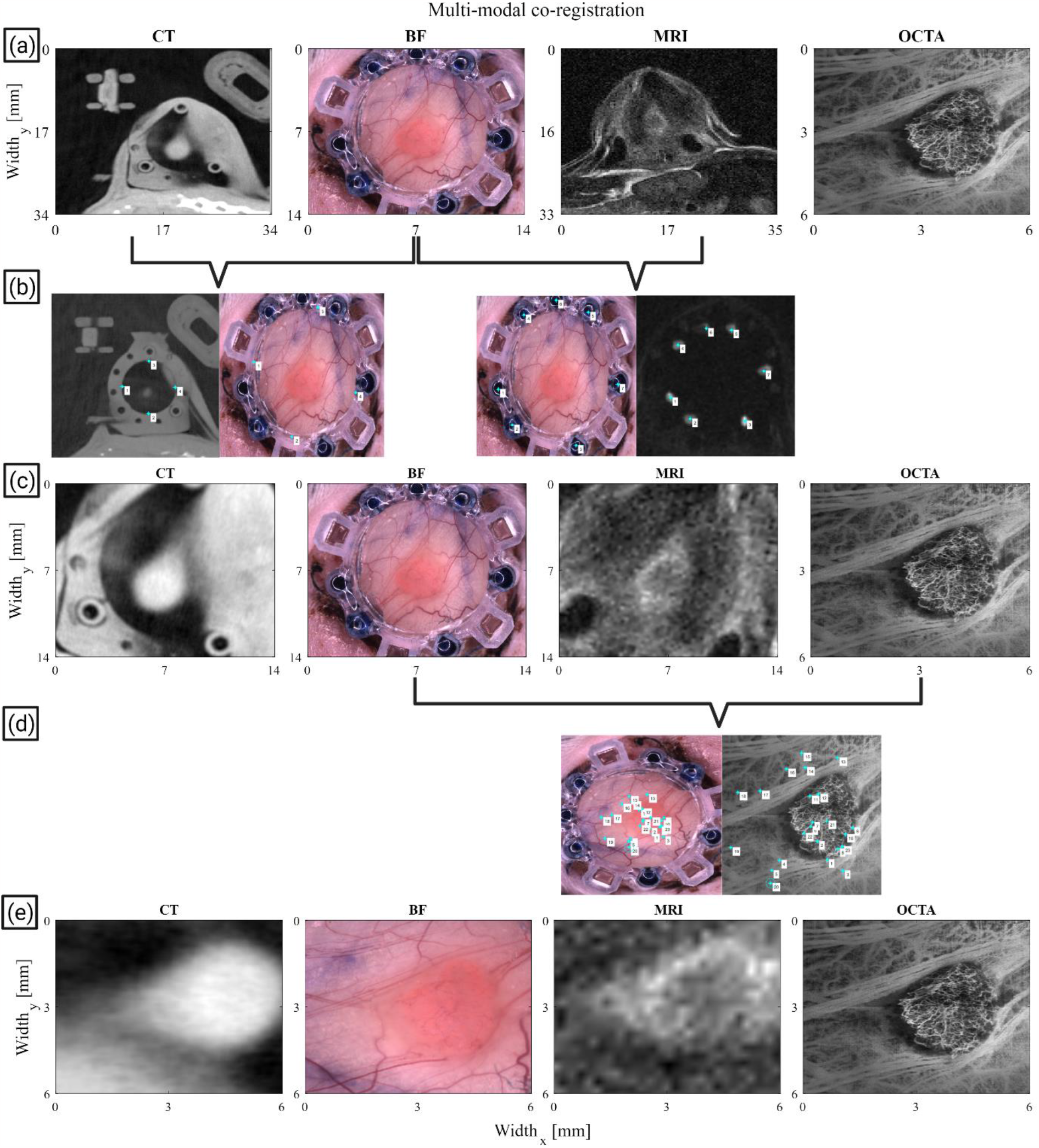
Visualization of the sequence of steps involved in multi-modal co-registration. (a) Representation of the original scale of each of the acquisitions; we note the significant lateral pixel scale and *field of view* (FOV) size difference between magnetic resonance imaging (MRI) (axial slice), computed tomography (CT) (axial slice), brightfield (BF) (and fluorescence, not shown), and svOCT (*en-face* average intensity projection) images of the same tumour in a mouse *dorsal skinfold window chamber* (DSWC). (b) First MRI is co-registered to the BF meso-modality (a middle ground resolution and FOV size modality between OCT and both MRI and CT) based on the visible water-dense and dyed wells (which appear as bright spots around the frame of the DSWC in the MRI acquisition). These fiducials are identified manually via MATLAB’s imregister built-in function (numbered white squares) to compute the required lateral similarity transform to co-register MRI to BF laterally only. Similarly, CT co-registration to BF relies on landmarking of the inner wall of the DSWC. (c) MRI and CT acquisitions are shown to be co-registered laterally to the BF acquisition in matching sized FOVs. (d) The BF meso-modality is being co-registered to the OCTA scan according to the visible vessels using another round of user-guided generation of the required similarity transform (highlighted on the brightfield and svOCT images accordingly in). (e) Finally, all modalities are co-registered to OCTA laterally. We note that at this stage all that remains is to complete axial alignment with respect to the window-air-tissue interface of the DSWC. Custom code for implementing this pipeline can be found in ^49^. All figures were prepared in MATLAB 2023a.

With all modalities aligned and superimposed, metrics computations from MRI (or other macro-modality) and OCT (or other micro-modality) can be performed over each 1 × 1 x 1 mm^3^ sub-VOI across the full tissue VOI, thus facilitating inter-modality correlations as further detailed in our previous study ^8^.

## 5 Conclusion and future work

In the presented work we demonstrate a novel DSWC design and imaging toolset enabling more robust 1) longitudinal, and 2) multi-modal pre-clinical imaging experiments. This pipeline has far-ranging potential applications for longitudinal studies in the biomedical sciences^1–13^. One such application is shedding light on the response kinetics of tumour xenografts to various therapeutics (e.g., chemotherapies, radiotherapies, etc.). Ongoing research is applying the pipeline described herein to improving our understanding of the impact of high dose per fraction radiation therapy on tumours and their stroma (e.g., blood microvasculature). This work aims to elucidate the long-standing debate on whether hypofractionated dose regimes rely on novel radiobiology and to develop a predictive model for the tumour response based on change to the stroma towards a future in adaptive radiation therapy.

This pipeline holds potential to reduce variability inter-timepoint and inter-specimen in notoriously complex *in-vivo* experiments. Beyond this, through improvements to multi-modal data acquisition and subsequent co-registration robustness, this pipeline may further facilitate studies correlating findings between pre-clinical modalities with clinically feasible analogues^7,9,49^. This simplification of translational studies may thus accelerate the transition from benchtop to bedside.

## Supporting information

Suppl 1 Evolution of the DSWC design and installation protocol

Suppl 2 DSWC Surgery Protocol

Suppl 3 Surgical stage construction

Suppl 4 Novel OCT-histology corregistration pipeline

Suppl 5 DSWC and toolset animation

Suppl 6 Automatic xy-microstage in practice

## 6 Acknowledgements

Thank you to Dr. Kathleen Ma and Dr. Anna Pietraszek, and staff most notably Mafe (Maria) Monroy at the University Health Network (UHN) Animal Resources Centre (ARC) for their support and feedback over the years of iteration of this pre-clinical model. Thank you as well to Dr. Warren Foltz and staff at the spatio-temporal targeting and amplification of radiation response (STTARR) facilities for their support in the operation of the small animal irradiator and scanner as well as imaging using the 7T MRI. Finally, many thanks to Dr. Timothy Samuel and Dr. Layla Pires for their initial support in preparing these animal models.

## Notes

https://drive.google.com/drive/folders/1BFr-JMqRf81g2N40xJZtYeX6mUx8XRaL?usp=sharing

https://www.thingiverse.com/naderallam/designs

https://github.com/nallam1/Batch-processing-multimodality-tumour-imaging

### Competing Interest Statement

The authors have declared no competing interest.

