## Supplementary material for "Low-Cost 3D-Printed Tools Towards Robust Longitudinal Multi-Modal Pre-Clinical Imaging": Suppl 1 Evolution of the DSWC design and installation protocol

Based on the criteria 3), 5) and 7), it was clear that additive manufacturing methods, such as the large field of plastic 3D printing would be highly suitable. In fact the polymer plastics used in 2 biggest forms of commercial 3D printing, fusion deposition modeling (FDM) and stereolithography apparatus (SLA), are largely made of organic molecular groups (hydrocarbon chains), thus addressing the issue of gradients in magnetic susceptibility, may be selected to have a range of material properties (biocompatibility^1–4^, glass transition point (temperature resistance)*, flexural modulus (bending resistance or stiffness), strength (load resistance at a given time) and durability (load resistance over time)^5^), as well as low cost & ease of use, enabling rapid prototyping.

* One of the preferred means for sterilizing equipment and tools (denaturing and destroying any residing potential harmful foreign biological material) envisioned for *in-vivo* are exposures to high-energy radiation (e.g., gamma radiation), or high temperatures (e.g., autoclaving)^6^. Methods of “cold” or chemical sterilization via oxidative or non-oxidative agents (such as the glutaraldehyde-based coagulant Cidex^®7^) are highly concentration and duration dependent and may therefore be more likely of performing incomplete sterilization. Autoclaves are most readily accessible, thus the chambers must be able to resist structural distortions resulting from prolonged exposures to high temperatures ~121°C for 15-30min as typical in autoclaving small tools.

To overview the evolution of the DSWC design:

1) Proof-of-concept model: At first a 1:1 scale imitation of the titanium DSWC from APJ trading was designed and FDM printed out of the widely-used and biologically inert polylactic acid (PLA) filament on a Prusa MK3S and Prusa Mini, including the nuts and bolts, with a rimmed opening in the frame intended to seat the 12mm borosilicate circular cover glass.

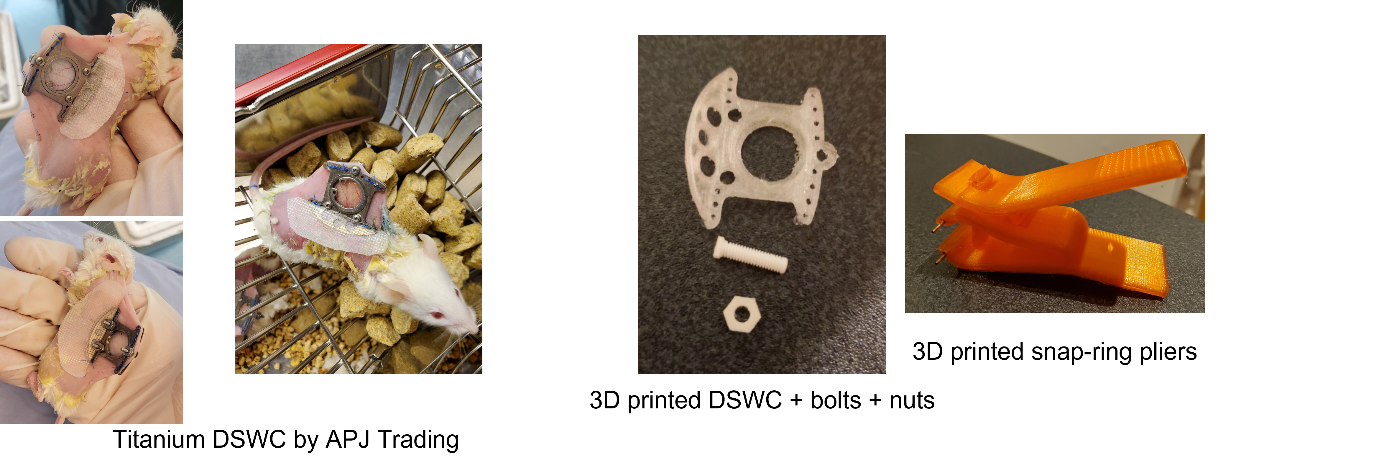

**Figure 1: The First plastic DSWC (taking inspiration from the titanium analog**). The glass inserted in the DSWC center is 12mm diameter for scale.

The preliminary issues encountered were:

I. Far too fragile for *in-vivo* use;
II. Not autoclavable (PLA without heat treatment has a glass transition point around 60°C).

2) Large-framed snap-ring model: Improving on these initial points to begin *in-vivo* testing, an improved design (more conducive to the durability of plastic relative to titanium), as well as more temperature-resistant materials and heat-treating techniques were investigated. These materials/techniques included polypropylene (PP) and oven-annealing PLA post-printing at 110°C for 30min (and allowing to gradually cool to room temperature over a period of hours^8,9^; this works in PLA because following rapid cool-down from 3D printing at ~210°C, the polymer structure is amorphous, however slow cooling results in a more organized and thus mechanically robust crystalline polymer structure). These different materials were representative of different stiffnesses: PLA is a hard (low flexural index) plastic and PP is a soft (high flexural index) plastic. All printing was performed on a smooth PEI sheet such that the surface in contact in tissue have less gaps for biofilm accumulation. It appeared that the softer PP, was easily gnawed by the rodents compromising the structural integrity of the chamber. While the harder annealed PLA DSWC had a far longer longevity (~2.5 weeks, nearing most other models lasting typically 3 weeks^10,11^, but far from permitting longitudinal observation a month or more post a ~1 - 2 week SBRT treatment) even though it was nearly double the mass of PP. The means for retaining the glass window in the window chamber frame depended on the metal snap-ring (as producing a similarly high-tensioned plastic version of the metal snap-ring would be very difficult and may not withstand the wear-and-tear of multiple sessions of snap-ring removal and insertion).

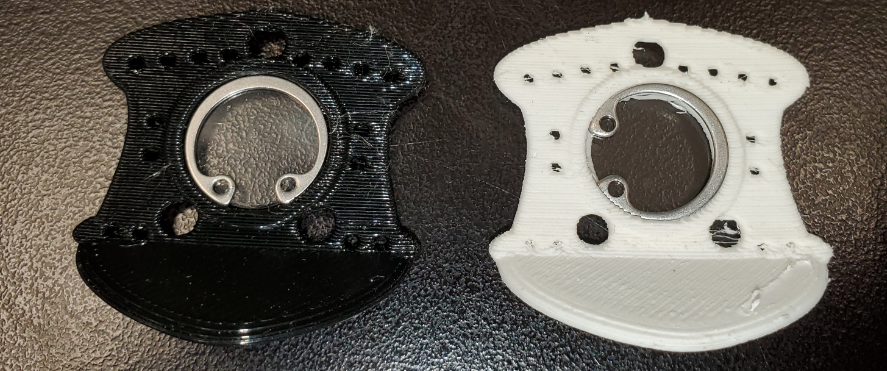

**Figure 2: Large-framed snap-ring model.(Left is PLA, right is Pegasus PP).** The glass inserted in the DSWC center is 12mm diameter for scale

It had several shortcomings, most notably:

III. No means of guidance for MRI-OCT co-registration; given the resolution difference of approximately 2 orders of magnitude, and differences in Field-of-view (FOV), no visual landmarks can be used to reliably co-register these modalities to one another.
IV. The snap-ring must be removed before each imaging session using a custom-designed snap-ring plier tool which applies pressure on the glass and tissue and may result in damaging of the glass or even air gaps and thus avenues for infection (in addition to perturbation of the tissue).
3) Large-framed O-ring retainer model: Solving the problem III., divots were introduced to the design at key locations on the DSWC, in which tear-gel (proton-dense) may be aliquoted prior to T2-weighted MR imaging (thus being captured in the FOV of MRI). These divots would also be optically visible and may be simply perceived in the FOV of the brightfield microscopy meso-modality. One attempt at solving problem IV. was the 3D-printing of retainer protrusions which the O-rings would be fitted underneath.

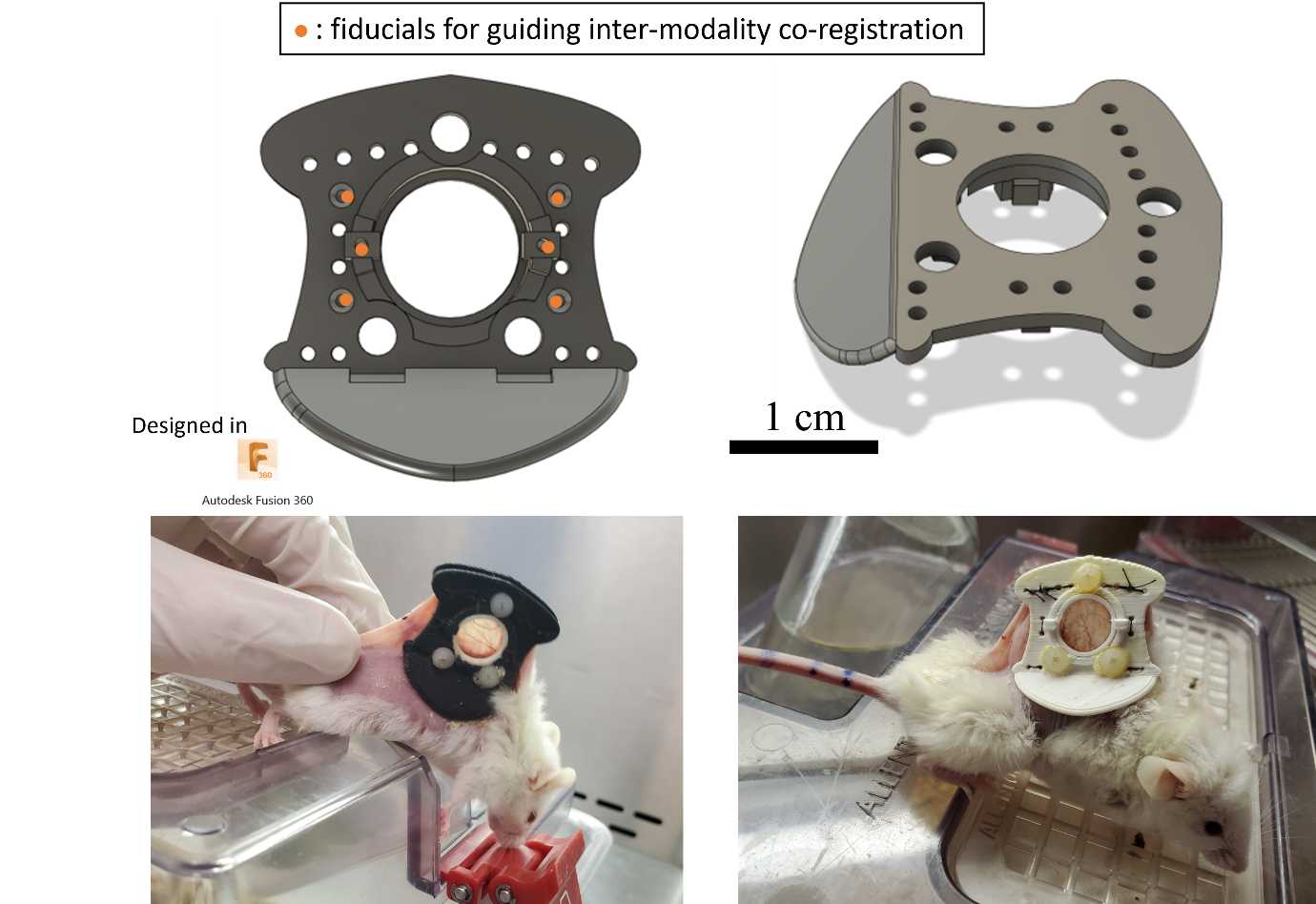

**Figure 3: Large-framed O-ring retainer model DSWC.** The glass inserted in the DSWC center is 12mm diameter for scale

The problems encountered:

V. O-rings may still be taken out too easily even by the mice, thus risking contamination of the DSWC (and gluing may result in undesired leakage onto the window). Additionally, the O-ring retainer setup was very bulky.
VI. The DSWC were exceedingly bulky and heavy, weighing approximately 3.5g (1g more than the titanium DSWC), approximately 15% of the mouse’s body weight. This would risk over-stretching of the skin and damage to the dorsal muscle, impacting mouse comfort, and compromising chamber longevity. Beyond this, temperature and humidity conditions in the extended dorsal skinfold are below typical bodily conditions in which pancreatic cancer for example would grow, larger chambers may likely only exacerbate the realism of this heterotopic model^12,13^.
VII. The process of annealing is very time consuming (and annealing results in structural distortions of the frame and shrinkage which were addressed via custom-designed PP and wooden moulds, in addition to over-compensations in the dimensions));
4) Compact glass-embedded model: To address V., the glass was embedded into the front-frame mid-print. I followed steps similar to other applications^14^, applying standard acrylonitrile glue to retain the light window in position once positioned, then resuming the print such that it is embedded in melted plastic. It may be noteworthy that a design Glass be glued from the backside of the front-frame was tested but was found to release from the frame overtime does representing a risk of contamination (justifying the need for an underlying rim supporting and “sandwiching” the glass). To properly address VI. the need for smaller and lighter DSWC frames, different material with higher strength and durability than PLA and high temperature resistance without the need for further treatment should be considered, thus simultaneously addressing VII. While the copolymer Acrylonitrile Butadiene Styrene (ABS) commonly used in injection-moulding (e.g., LEGO^®^) is another commonly available material (among the first developed for FDM 3D printing) known for its high strength and durability, its potential toxicity during printing, its glass transition point just short of the standard autoclave temperature, and solubility in acetone along with acrylonitrile glue (acetone is used in the post-processing of chambers to clean evaporated glue residue) left us considering alternatives. I finally decided on the high strength and durability blend consisting of another copolymer, nylon, reinforced with carbon-fiber (CF) (heavy-duty CF-nylon filament^®^, Filaments Inc, Toronto, ON, CA), with heat resistance up to ~167°C, with lower density than pure nylon. Thus using this material, more compact (by ~5mm) prints lighter than the titanium DSWC are possible (reaching ~2.1g).

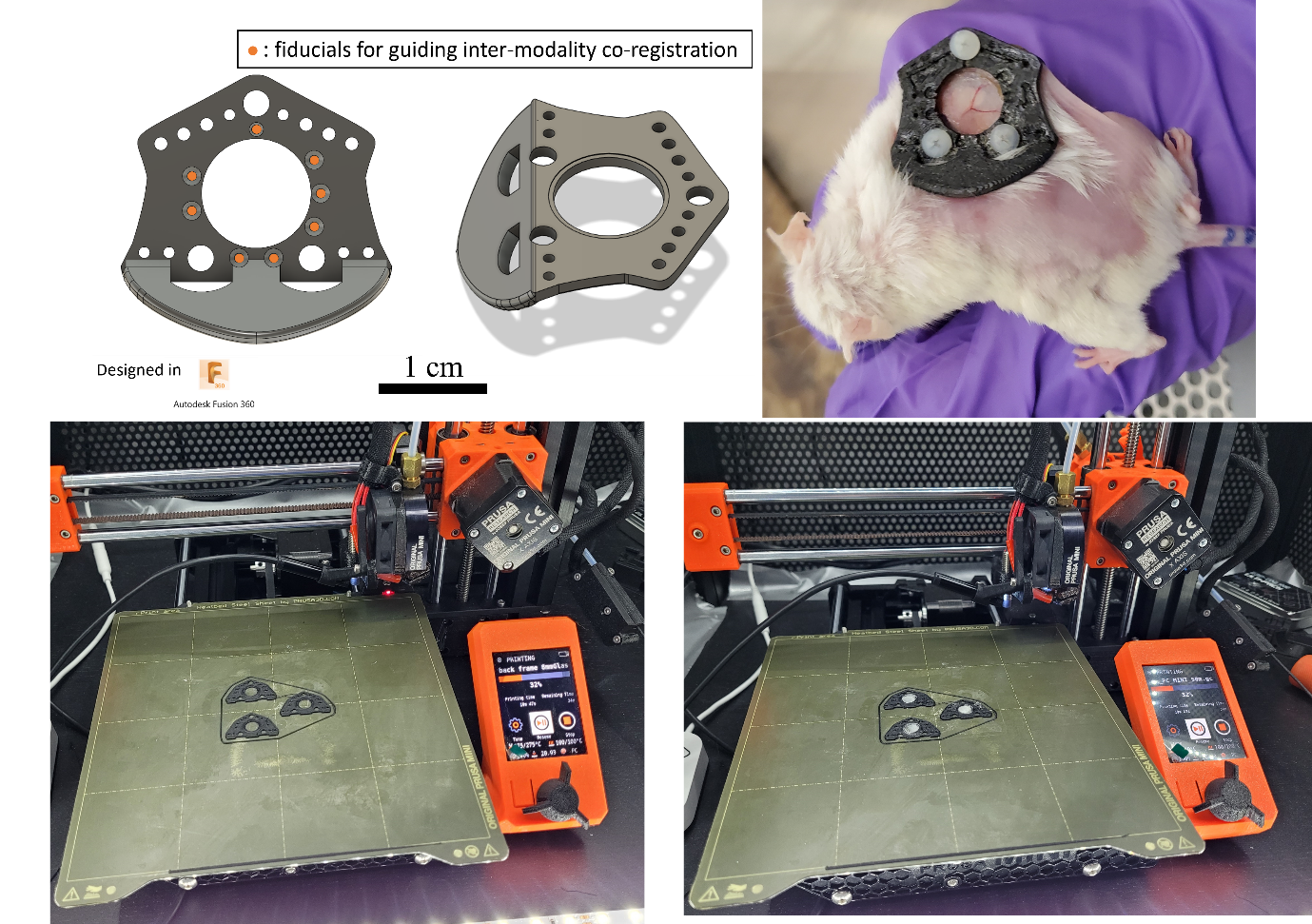

Figure 4: Compact glass-embedded model. The top shows details pertaining to the model. The bottom shows the steps involved in glass embedding. The glass inserted in the DSWC center is 12mm diameter for scale

The main issues however were:

VIII. At sacrifice, 3-5 weeks post DSWC installation, in all models to date, mice would exhibit signs of necrosis (e.g., discoloration) in the tissue directly sandwiched by the DSWC.
IX. At the time, I had not conceived of a design for screws which may be FDM 3D printed which may be sufficiently durable all the while maintaining a small diameter on the order of 1mm with threading as in titanium DSWC. Therefore, the best solution appeared to be purchasing M2.5 nylon nuts and bolts typically used in applications requiring thermal and water resistance^15^. These were secured required acrylonitrile glue (Gorilla glue^®^ worked better than Krazy glue^®^ on CF-nylon). Such screws required large and potentially traumatic incisions in the mouse skin, which may compromise DSWC longevity.
5) Standardized compact glass-embedded model: Addressing problems VIII. and IX., CF-nylon spacers (~1.6-2 mm thick, empirically determined to work best for NRG mice, thus standardizing the inter-frame spacing) were designed and printed along with CF-nylon 1mm thick bolts (rather than screws) with a bevelled tip (to pass through the skin less traumatically). These bolts pass through the front-frame followed by the spacer, and the back-frame before being fixed in position with a cap/stopper, I call “bolt sheath” which also play the role of protecting the mouse against the sharp tipped bolt (which are not simply cut as these bolts help in fixing the DSWC to the imaging bed during imaging). These are CF-nylon printed alternatives to screw-on nuts which tightly fit onto the bolts (thus avoiding the issue of printing threads).

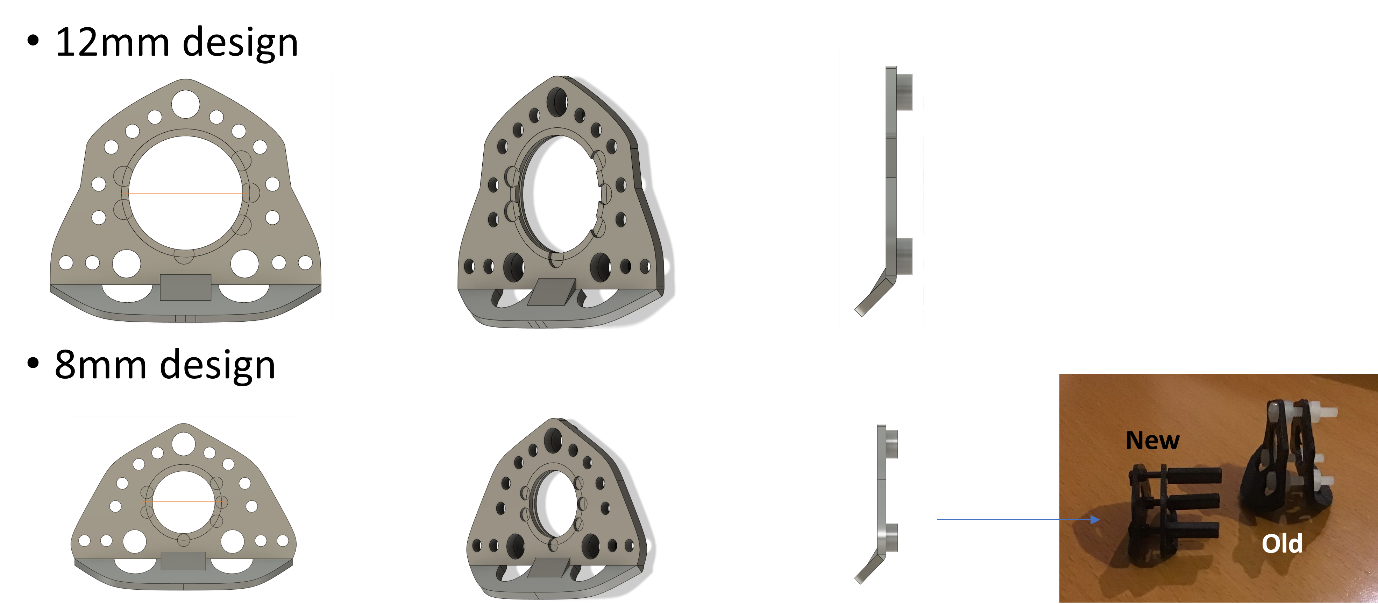

**Figure 5: Standardized compact glass-embedded model.** We note that the latest designs are more compact and now include spacers, and use far smaller bolts.

With these improvements implemented improving model longevity (i.e., by minimizing sources of animal distress) and accuracy (e.g., reducing impact of phenomena observed based on variability in tissue perfusion), further issues may be addressed:

X. Over the period of weeks post-DSWC installation, as skin elasticity dimishes^11^, tissue drift in the DSWC may occur as well as lateral tilt in the DSWC^10^.
XI. Weeks following window chamber installation, transudate (~transparent/translucid) and exudate (highly turbid) are released, crucially obscuring the FOV to optical microscopies. This may be a result of infection or a result of elevated cytokine activity as an inflammatory response to the skin wound, characterising all WC installations, towards repair^16^. It has been noticed that if an air gap is present over the exposed panniculus carnosus muscle tissue, exudate formation is exacerbated (likely due to osmotic pressures driving fluid outside the tissue).
6) Glass-embedded semi under-skin model: In the 6^th^ design iteration, it was clear that to address X., the DSWC must be made lighter and more compact, reducing the skin tension and sustained load applying torque to the skin fold as much as possible, all the while improving how tightly the tissue is fixed in position, having increased support to help the DSWC maintain a perpendicular orientation to the mouse’s back (Previously achieved via 45^o^ flaps at the bottom of the DSWC frames), and maintaining space for the fiducial markers in the resulting minimalist design. The front-frame thus became a simple disk with embedded glass, with suture holes along its circumference and divots for the fiducials, while the back-frame, became more compact with an insert for the “hula-clip” support resting on the mouse’s back (thus applying minimal torque to the skinfold) at 50 mg. Addressing XI., in an attempt to reduce chances of air gap between the tissue and the window, mitigate sources of infection and improve tissue fixation in the DSWC, the 17 mm diameter and 1.7 mm thick front-frame was inserted underneath the skin, similar to the installation protocol of abdominal WC, thus doubling the number of transdermal fixation points per suture, in addition to further sealing the DSWC from foreign bodies via a biocompatible acrylonitrile glue known as Vetbond^®^. At this stage, the DSWC works very well, already (~1 g) lighter than titanium DSWC, and having a potential longevity of 1-1.5 months post-installation.

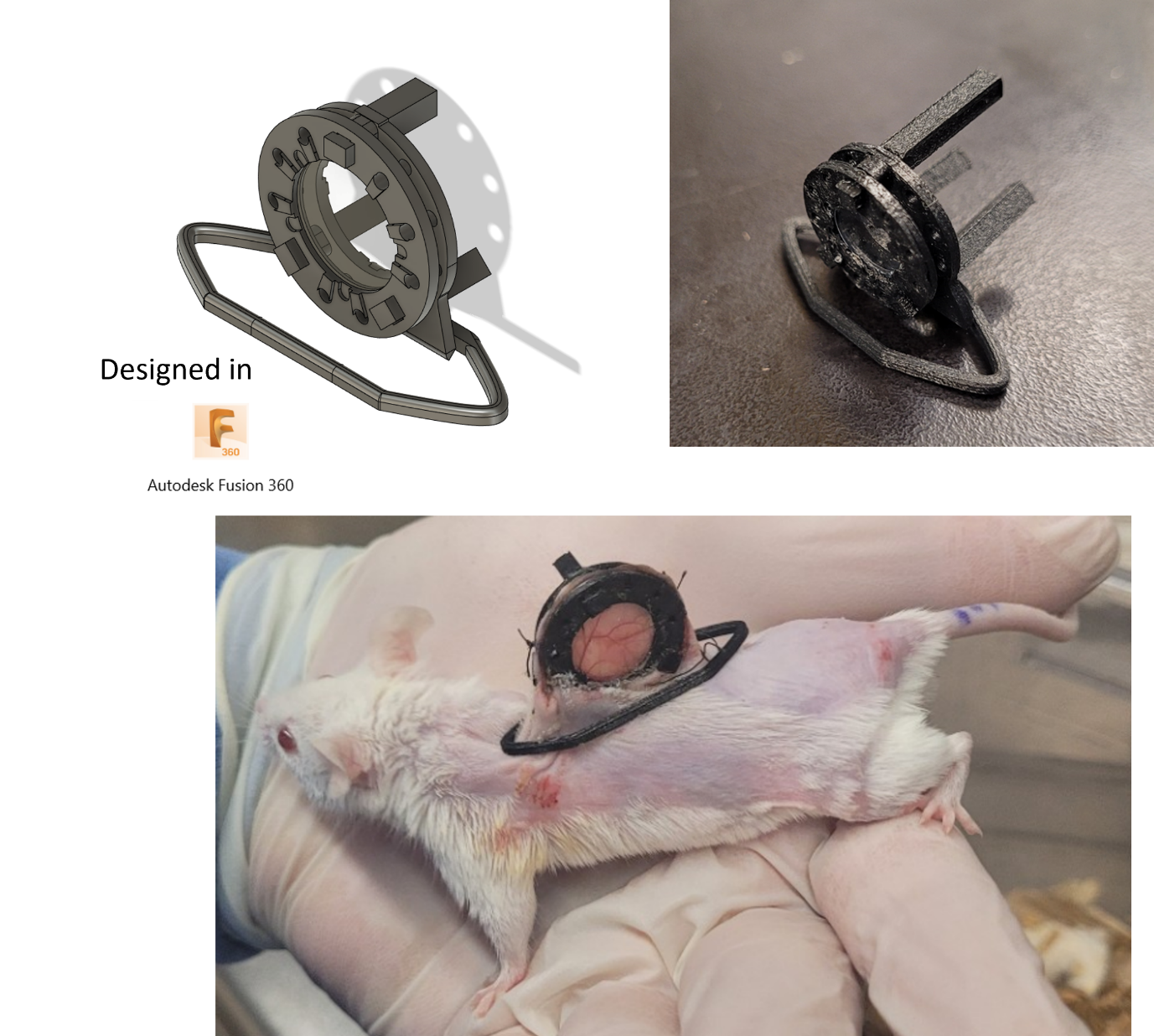

**Figure 6: Glass-embedded semi under-skin model**. The glass inserted in the DSWC center is 12mm diameter for scale.

However clear issues remain which may lead to further improvements:

XII. DSWC preparation is a slow and complex process: Glass embedding was a meticulous (and painful!) manual process at the risk of hands sticking to the 100°C printing bed (as tweezer were found to be difficult to manipulate the small glass cover-slips into the low-tolerance diameter frame opening onto the rim). The success rate in embedding was also close to only 50%, improving marginally with experience, due to small artifacts in the printing or imprecise positioning in the glass resulting in its fracturing as the print resumes. The post-processing required consisted of the removal of stringing artifacts and submersion in 99% acetone followed by light polishing of glass with lint-free optics wipes (e.g., Kimwipes^®^) and rinsing with water (tap-water is satisfactory).
XIII. Over the course of the weeks of treatment and imaging and even during surgery, the thin glass window could be fractured, becoming a risk for infection and potentially compromising the imaged FOV.
XIV. Dirt accumulation on the DSWC window also becomes problematic over time given that we are performing optical imaging. This is exacerbated by the need to remove snag hazards from inside the mouse cage, including the grading which usually holds the mouse food. While cleaning with acetone, alcohol or other typical glass polishing agents helps, there is sometimes difficult to reach accumulation at the inner rims of the frame, and pressure on the glass during cleaning may result in its fracturing.
7) Back to O-rings: Addressing issues XII., XIII and XIV. was far from straightforward but linked. Toward mitigating XIV., various animal care measures have been improved including the design a mouse feeder, along with regular cleaning of the cage, as well as a change in DSWC cleaning protocol, whereas rather than relying solely on wiping the window via polishing agent prior to each imaging session, a hand-powered air-blower is used (see Figure 7). One considered solution toward solving XIII. and XIV., was the design of a protective belt, however the only problem was that it was far too bulky and FDM printing could not practically implement this solution with protrusions (given the anisotropy of parts durability as a function of printing orientation). The seemingly best solution to these 3 problems (XIV in part) appeared to be to revisit the O-ring concept, permitting the glass to be replaceable as needed, however reducing dimensional tolerance such that it would only be possible to insert/remove the glass and O-ring with great care via forceps.

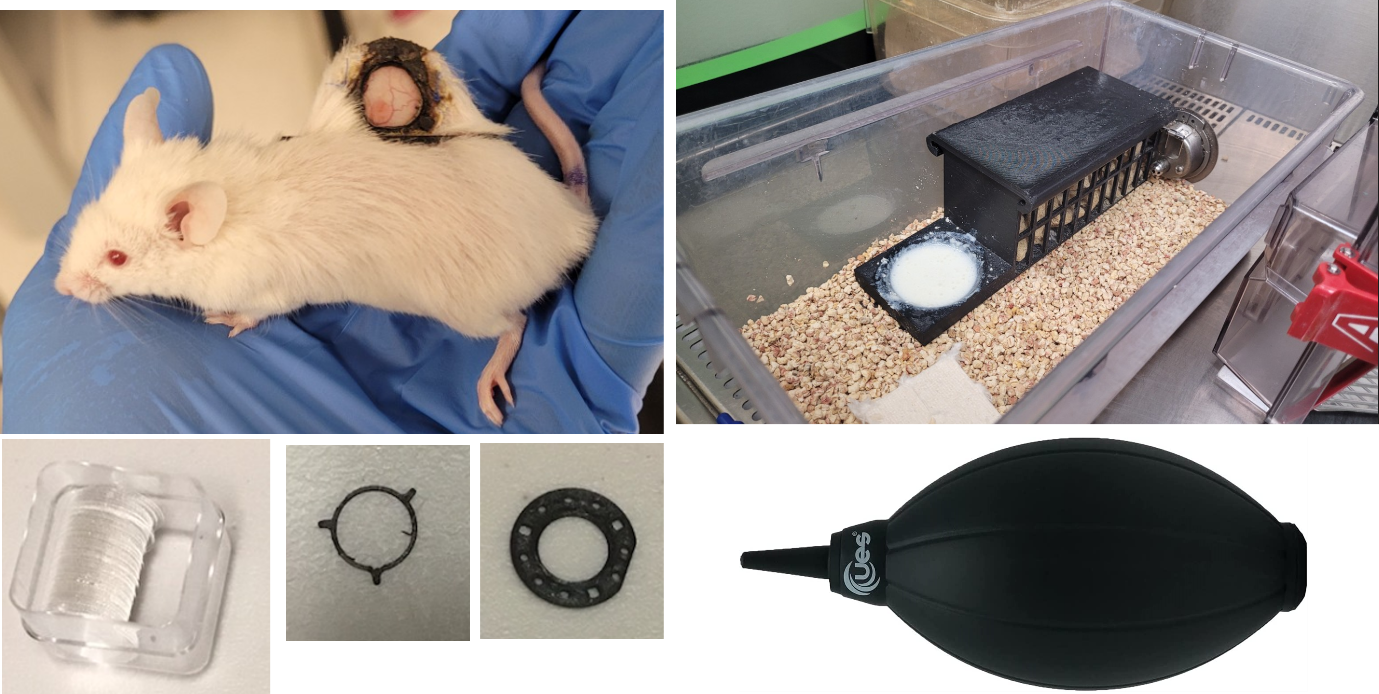

**Figure 7:Back to O-rings: Semi-under skin with O-ring model** (left). On the right are some of the aforementioned solutions to the dirtying of the DSWC (air blower and mouse feeder).The window is 12 mm for scale.

While this solution did work as expected, it had several problems:

XV. It worked too well: it was often too challenging even for the experimenter to insert/remove the glass and O-ring without fracturing a few windows and having to replace them, which was not only time consuming but also came at the risk of damaging the exposed panniculus carnosus muscle tissue.
XVI. Tear-gel cannot be precisely nor repeatably manually aliquoted in the divots;
XVII. During surgery, it is often challenging to center the tumour in the DSWC and the issue may be a result of inaccurate positioning during tumour inoculation.
XVIII. Additionally, the use of injectables anesthesia (e.g., ketamine + xylazine cocktails) during surgery, have largely unpredictable durations of action (thus leading to potential distress in the mice or worse, attrition), unlike inhalants, (e.g., isoflurane), which are more dynamic in action, remaining functional only as long as the specimen is exposed, in a depth of anesthesia a direct function of the isoflurane concentration and flow rate. Overdosing is also easier to overcome with isoflurane over ketamine/xylazine, as it would simply be a matter of providing the specimen with an “oxygen flush” (removing the source of isoflurane and, optionally but recommended, delivering pure oxygen to the animal until they become ambulatory).
8) Full resin with permanent fiducials model: Already from XV., it was clear that reverting to permanent window designs maybe best to reduce the risk of damaging the very tissue we are trying to image. Different thickness and size glass coverslips were tested, and though reducing the size did help in reducing the risk of fractures, this solution would be a compromise to our potential imaging FOV. A radically different solution was contemplated: given the durability and flexural modulus of plastics, why not try to 3D print the window rather than rely on glass? While the idea of 3D printed optics is not new, being an important part of the rapidly evolving field of integrated photonics, these tend to use complex printers (e.g., direct-ink writing, two-photon laser printing, drop-on-demand inkjet printing)^17–20^; extrusion based printers like the commercially available FDM printers may have too significant variability in print density (as a function of the material used and flow-rate variations) leaving imperfections such as air bubbles, surface roughness (not only as a function of the alignment and type of substrate used) and deviations from the intended design^20^. Fortunately for our intended purpose, we are not designing complex optics; we simply require a window, a smooth homogeneous slab of optically and NIR (~1300 nm) transparent material of ~constant thickness and index of refraction (thus minimizing scattering and wavefront aberrations). I had initially attempted to use FDM printing with various materials (e.g., natural PLA (without dye additives), different formulations of the copolyester polyethylene terephthalate glycol (PETG) including Colorfabb Clear HT PETG^®^ (with high temperature resistance), polyvinyl butyral (PVB, an alcohol soluble resin), etc.) according to a range of printing parameters and applying smoothing solvents where applicable^21–23^. Most achieve the greatest results in transparent 3D printing via the logistically more complex SLA 3D printing, the 2^nd^ biggest form of commercially available 3D printer. Thus, building on my home setup, I began to use the Elegoo Mars 3 masked SLA (MSLA) Here rather than relying on the deposition of melted resin by a mobile extruder line by line onto a substrate (FDM 3D printing), the substrate is periodically lowered into and raised from a vat of photoreactive resin, polymerized layer-by-layer according to the pattern illuminated (at ~405 nm) by a screen at the bottom of the vat. MSLA printing can achieve significantly higher resolution parts (down to $10 \mu m$ axially (controllable according to the height of the substrate) and $35 \mu m$ laterally (fixed by the pixel count of the MSLA printer), an order of magnitude or more over FDM printing which is limited by the belt, linear motor, and nozzle diameter used). Thus, a novel more compact DSWC model was printed in Formlabs BioMed Clear Resin (formlabs inc.), a USP Class VI certified material designed for long-term skin and mucosal membrane contact, at $50 \mu m$ axially according to their recommended printing, curing and post-curing protocol^24^. Along with printing the DSWC and window entirely out of resin, the attempted solution to problem XVI. was the design of ~.8 mm diameter and ~1.6 mm deep cylindrical wells to be filled with a proton dense fluid which may also be easily captured via brightfield imaging, similar to the way certain radiology departments may stick vitamin E pills onto patients as a low-cost fiducial^25^. Thus, after considering many of the successful low-cost materials tested by Izatt et al.^25^ (i.e., fish oil, and vitamin D), finding these were difficult to optically capture (and being hydrophobic, food coloring dye therefore does not mix), the wells were finally filled with a mixture of tear-gel with a few drops of blue food-dye, via insulin syringe. It is crucial that these wells be tightly sealed such that the fiducials do not dry over time, rendering them obsolete in longitudinal MRI – OCT co-registration experiments. Thus, after post-processing the prints and then filling the wells with the fiducial contrast agent, these were manually sealed with a small drop of resin photopolymerized via hand-held UV-lamp. The seal-quality was assessed via submersion of the completed DSWC in a tub of water and watching for the diffusion of the dyed solution from the wells. Finally addressing problems XVII. and XVIII., a surgical stage and “surgical guide” was designed to enable this full DSWC installation procedure to be performed under isoflurane anesthesia. While several design iterations were built and tested to different extents (e.g., 3D-printed loc-line tube adapter piece, over-head surgical stage, rotatable heated stage), the best design to date consists of a custom-designed heated mouse-bed with isoflurane inlet tubes fitted onto a third-arm soldering stage (Kotto^®^)^26^ meant to help manipulate and fix the “surgical guide”. The latter consists of a “c-clamp”, to which the mouse dorsal skinfold may be stitched, with adaptor hole to tightly fit the DSWC back-frame; previously the DSWC back-frame would have to be positioned and held against the dorsal skinfold with one hand while the other hand is used to attempt to precisely mark the bolt holes and central skin flap to be removed. Given the skin elasticity of the mice, attempting this process of accurately and precisely marking the DSWC placement with tumour centered (as well as 25 days prior, during the determination of the ideal position for tumour inoculation (the position most-likely to enable tumour growth at a location conducive to DSWC placement with minimal animal distress and for most upright DSWC orientation upon installation)) was largely guess-work prior to the development of the “surgical guide” helping to manipulate the skinfold (refer to **Error! Reference source not found.** on page **Error! Bookmark not defined.**).

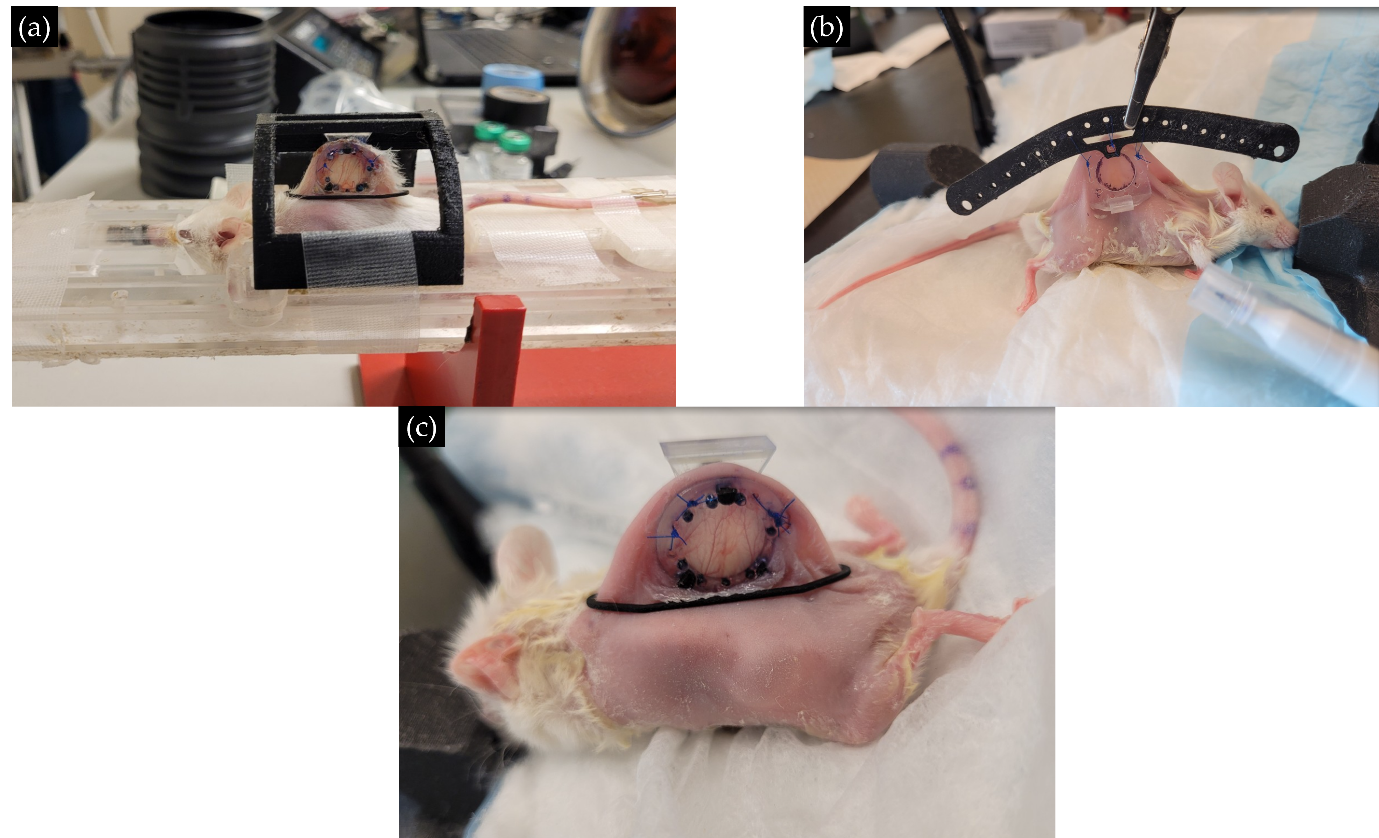

Figure 8: Full resin with permanent fiducials model. (a) Mouse lying in MRI bore with backframe hooked into MRI-restrainer for guided alignment and reduced motion artifacts. (b) Surgical guide held by 3^rd^ hand of surgical stage during surgery. (c) The mouse post-surgery. The window is ~11mm for scale.

Nevertheless, as anticipated from such a radical change in approach, some issues were encountered:

XIX. The fiducial markers resin-seals would often be broken by the activities of the mice, leading to drying of the fiducial wells.
XX. Though scratching of the plastic window was not observed, it is well known that plastic is far less scratch resistant than glass. It is therefore important to protect the DSWC from this, along with dirt from the animal activities while in the cage (as in problem XIV. which had not been fully addressed because of the limitations of FDM 3D-printing).
XXI. Given that the substrate properties (leveling and texture) and vat tension impact the quality of print, though the parts obtained are highly transparent for what may be expected of 3D-printed plastic, texturing artifacts (grid-like pattern) appear at the surface of the window resulting in diffuse scattering from the window surface, and thus blurring.
9) Full resin with replaceable fiducials: Contemplating XIX., I realized that the fiducial markers had no need to be permanently installed on the DSWC. Therefore a replaceable fiducial marker holder may be mounted on the frame during brightfield/fluorescence as well as MR imaging and removed upon completing the imaging session… but how to ensure consistent positioning of the replaceable fiducials? The front-frame was designed with small thin protrusions to which the replaceable fiducials (and any mount with the correct dimensional tolerances) may be tightly fit; this is possible as the mechanical properties of SLA printed parts are isotropic by the nature of the technique. This same solution may be used for mounting the DSWC “window shield” (front-frame cap for protecting the frame during mouse daily activities in the cage), towards problem XX. Finally, towards solving XXI., various techniques were considered to help replicate the optical clarity of glass with MSLA printed resin. These included heating on smooth substrate, mechanical polishing, solvents (e.g., isopropyl alcohol, acetone, ethyl acetate), and coatings (clear-varnishes and clear spray-coats)^20,22,23^ . While both nail-polish varnish and clear spray-coats were equally the most effective at achieving high optical clarity, it is also crucial that the selected methodology be easily repeatable, which was not the case for nail-polish varnish which had to be diluted in acetone to an empirically determined concentration and carefully manually spread on the window with minimal pressure to leave no impressions upon drying. Thus it was found that spraying at a distance of ~5-10 cm with USC SprayMax 2K Glamour High Gloss Aerosol Clear^27^ in a fume-hood and leaving to air-dry in the fume-hood worked very well, quickly, and consistently. As this spray coat is not designed to be biocompatible, the backside of the front-frame was masked by adhering it to off-the-shelf polyvinyl chloride (PVC) tape (packing tape) during the spray-coat process.

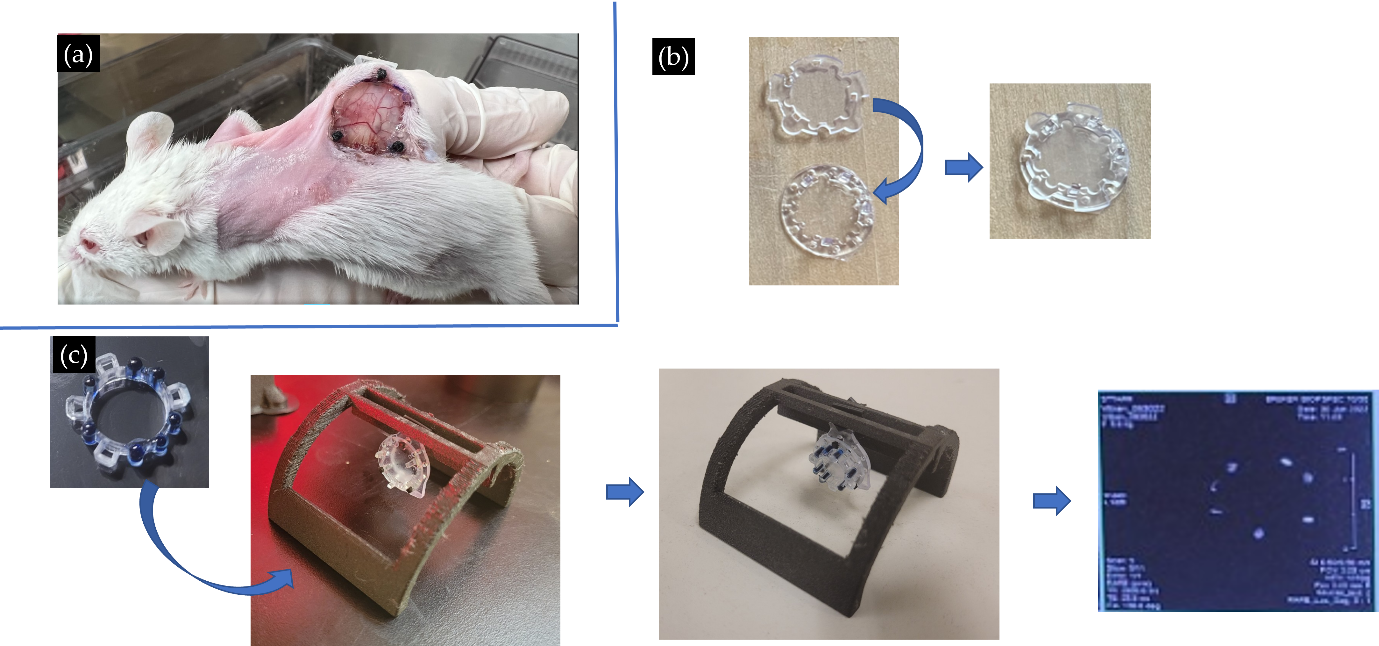

**Figure 9: Full resin with replaceable fiducials.**(a) DSWC alone (notice the hula-clip is not always necessary to be placed (as it may have the effect of becoming a nuisance to the mouse if placed in such a way as to obstruct motion). (b)Protective cover for the DSWC front-frame. (c) Replaceable fiducials shown here being placed on a front-frame already positioned with the back frame in the MRI restrainer piece, a T2 scan shows that indeed the fiducials do appear of high contrast (signifying no (or very little) leak is occurring which will not be exacerbated by potential damage through the mouse’s daily activities.

The final model presented in this manuscript omits the transdermally inserted bolts, to further improve model longevity^10^. All-in-all, after multiple design iterations and tests, to our best knowledge, this design addresses all previously raised issues and meets the enumerated constraints in **section 3.2** of the manuscript.
