## Supplementary material for "Low-Cost 3D-Printed Tools Towards Robust Longitudinal Multi-Modal Pre-Clinical Imaging": Suppl 2 DSWC Surgery Protocol

Suppl. 2: Dorsal Skinfold Window Chamber (DSWC) Surgery

Materials:

| Drugs:   - Anesthetics (isoflurane or ketamine/xylazine mix) - Analgesics (buprenorphine) - NSAID (Metacam)   Other:   - 7.5% Betadine scrub solution - 10% Betadine solution - Cotton gauze or sterile paper towels - Electric heating pad - Electric razor - Hair removal lotion - 70% isopropyl alcohol - Induction box (3D-printed) - IR gun thermometer (optional) - Saline solution - Scale - Soldering iron with 1 mm tip - Sterilizing beads Tear gel - Surgical guide - Surgical marker - Surgical Tape - 3 Syringes (with needles) - Tissue adhesive (Vetbond) - Vernier callipers (if spacers are home-printed) | Tools:   - Forceps - Needle driver - Sutures - Suture scissors - Surgical scissors   Apparatuses:   - 3 window chamber pieces (Formlabs Biomed clear resin printed): Front frame, back frame, and chamber protector. - Hula clip - Surgical stage with third-hand + surgical guide |
| --- | --- |

Prior to date of procedure:

1) Ensure all supplies available (status of: Canadian Drug exemption, AUP, etc.)
   a. Aliquot drugs (Metacam (5mg/kg, 0.5mg/mL))
   b. Submit technical request of SR buprenorphine analgesia ahead of time, including mass of mice. Do not forget to mark the cage with a cage card, flagging the request, including mass of mice, date of procedure, lab contact and AUP.
2) Book surgical Biological Safety cabinet (BSC) or microsurgery suite well in advance.
3) Book anesthesia machine well in advance.
4) Ensure the suture holes of DSWC frames are fully clear. In case of obstruction, a 1 mm tip soldering iron can be used.
5) Dry autoclave metal and 3D-printed tools and apparatuses (resin-printed materials are sterilized via 5min submersion in 70% isopropyl alcohol.

Day of procedure preparation:

1) Carry all materials in toolbox and on trolley.
2) Once in vivarium:
   a. Prepare new mouse cages without mouse toys or top grating of cage (post-surgery card can be prepared while awaiting mice to regain consciousness), add sterile paper towel, 3D printed mouse feeder and if possible KMR mix dissolved in autoclaved water (see **Figure 1**).

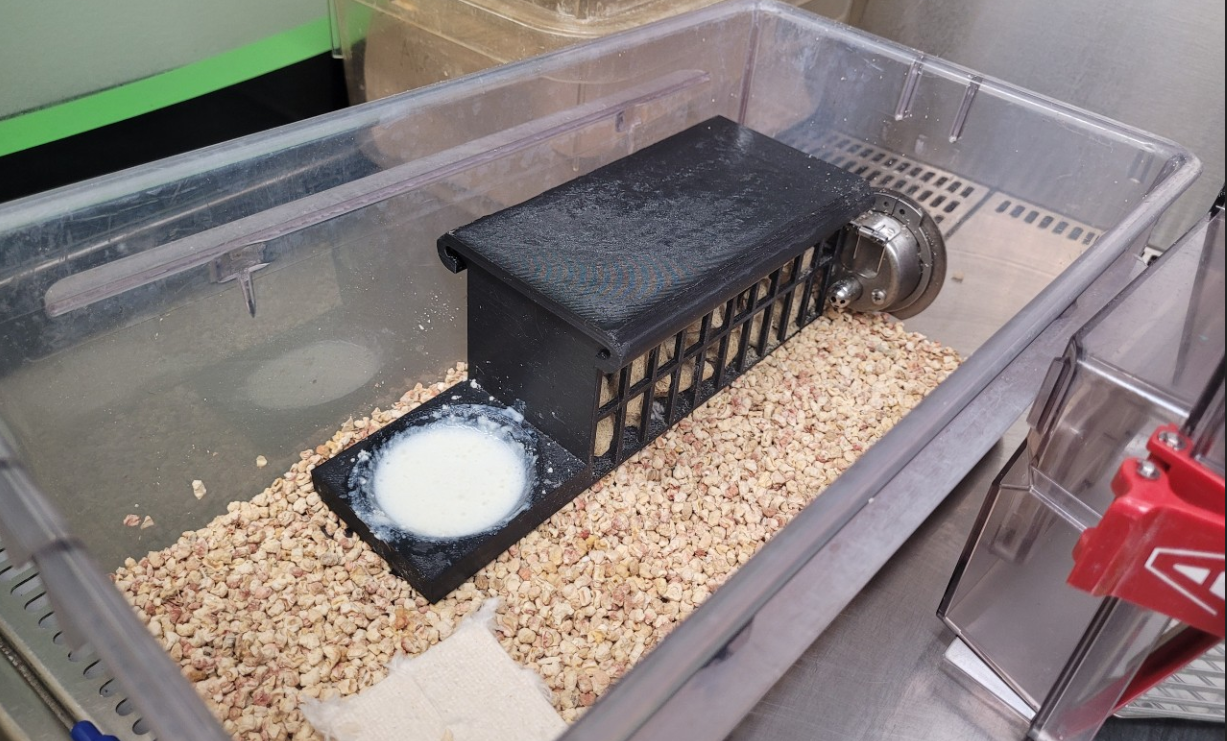

Figure 1: Clean cage prepared for post-operative recovery. The clean cage includes no snag-hazard objects, only the 3D-printed mouse feeder with drinking bowl. 5% irradiated mouse food pellets and KMR mix is given to the mice

b. Remove given cages of mice from vivarium individually ventilated cage (IVC) racks and drape with ARC gown.

3) Sterilize surgical table (Quarticide then Virox left to settle for 5min)
4) Set up two surgical pads: one above the surgical stage for operation and the other above an electric heating pad for pre- and post-surgical recovery.
5) Weigh mice and prepare and label 3 syringes (Metacam NSAID). You may use tape to help mark the syringe.
6) Submerge all Formlabs Biomed clear resin printed apparatuses in 70% isopropyl alcohol for 5min at least to disinfect (in addition to autoclave sterilization). See **Figure** 2a)

***Procedure:***

1) Anesthetize the mouse with 5% isoflurane in the induction box.
2) While waiting for the mouse to go unconscious, apply surgical tape on the hula hoop support part of the window chamber to ensure it sits firmly on the mouse without irritating the skin.
3) Apply tear gel every 20min, as necessary.
4) Shave hair from surgical site with electric razor. Continue to remove residual hair with hair removal lotion, as in **Figure 2b)**.
5) Within 30s-1min, wipe off lotion thoroughly with wet paper towels (water).
6) Clean the shaved area with 7.5% betadine followed by 70% isopropyl alcohol and 10% betadine. You may repeat this process for a total of three times.
7) Allow the mouse to dry-off.
8) Transfer the mouse to the sterile working area (ontop of Third-arm surgical stage), as in **Figure 2c)**.

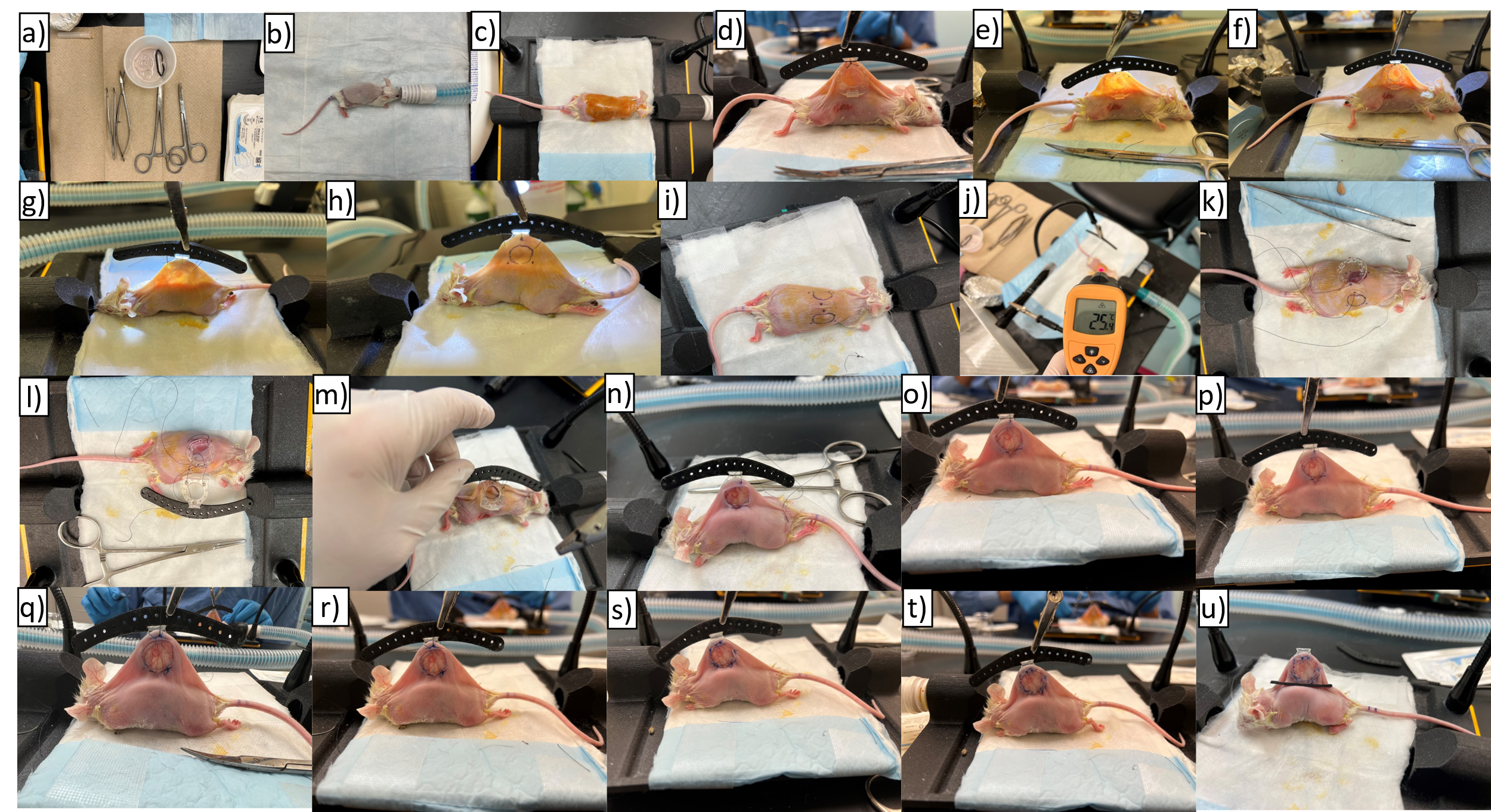

**Figure** 2: Various steps of DSWC installation. Steps: a) Preparation of sterilized surgical tools and DSWC components; b) Shaving mouse under ~1-2% isoflurane at 0.5 L/min in pure O_2_ on sterilized heated underpad; c) Sterilization of skin and transfer to heated surgical stage; d) Initial landmarking via surgical guide tool; e)-h) Transillumination of suspended skinfold to guide suturing and tissue resection for front frame insertion; i) Release of surgical guide to commence surgical attachment of full DSWC; j) IR gun reading confirms mouse skin at 24-30°C ; k) Removal of skin flap to expose panniculus carnosus, and initial suture to front frame (initialization of criss-cross temporary suture pattern outlined in step 19)); l) Intermediate steps of criss-cross suture pattern securing back and front frame; m)-n) Completed criss-cross suture pattern (back and front view respectively) and securing to surgical guide tool; o-s) Suspending surgical guide via third-arm clip and initiation of permanent sutures in order specified in **Figure 3**; t) Complete installation of DSWC including addition of chamber protector and removal of temporary sutures, and u) Removal of surgical guide and addition of hula hoop support.

9) Push the back frame through the surgical guide.
10) Establish a landmark for proper DSWC placement by using the window chamber back frame (connected to the surgical guide) on the side of skin containing the target tissue to be imaged (e.g., if the mouse is tumour bearing, position the back frame against the side with the tumour) and pull skin up together with one hand such that:
    a. The top of the skin fold clears the top bolt hole
    b. The bottom of the window chamber frame is sufficiently high above the mouse back to allow for the frame to rest comfortably on the mouse’s back without affecting the mouse’s breathing or locomotion.
    c. If mouse is tumour bearing, pull skin up such that tumour (target) is slightly (~2mm) offset lower than final desired location (e.g., for the tumour ­­to be ideally centered in the chamber, aim to have tumour ~2mm below center). *N.B.* It has been empirically noticed that leaving supplemental loose skin from the top improves the longevity of the chamber as it reduces the tension on the skin.

See **Figure 2d)**.

11) Ensure through a toe pinch that the mouse is in the surgical plane of anesthesia,

Suture the top hole of the back frame (BF_T_) through the skin to the central hole of the surgical guide. See **Figure 2d)**.

12) Suture nearest or second nearest suture hole of the back DSWC frame through the skin to the surgical guide such that the tissue in the back frame is under sufficient tension for drawing the contour guiding the next steps of DSWC installation. See **Figure 2f)**.
13) Suspend the surgical guide via the surgical stage third-arm, as in **Figure 2g)**.
14) Using a surgical marker draw a circle along the inner edge of the back frame and dots on the tissue behind the spacer holes, as in **Figure 2h)**. *N.B.* The spacer markings are important to guide the complex “criss-cross suture” pattern described in step 20).
15) Carefully rotate the surgical stage without disturbing the suspended surgical guide and tissue draw an identical circle and three dots corresponding to the spacer holes opposite to that drawn in step 14). Using the suspended flashlight may help in tracing these accurately (and potentially seeing the DSWC frame across the skinfold), see **Figure 2i)**. One may also choose to guide the marking of the front frame side by pushing needles through the back frame and skinfold from the backside. *N.B.* The spacer markings are important to guide the complex “criss-cross suture” pattern described in step 20). *OPTIONAL:* Draw a straight line along the left to right sides diameter of this **2^nd^** drawn circle extending it by an additional 1mm from either side.
16) Remove the sutures connecting the surgical guide and backside frame, see **Figure 2j)**.
17) Use forceps to pick up the skin at the center of the **2^nd^** drawn circle (from step 16) and cut the circumference of this drawn circle to remove the forward-facing portion of the skin, leaving all the vessels on the opposite skin side intact. Use forceps to remove any remaining loose fascia, see **Figure 2k)**.
18) *OPTIONAL:* Cut along the extended diameter line from step 16.
19) Place the front frame and begin the criss-cross temporary suture pattern:

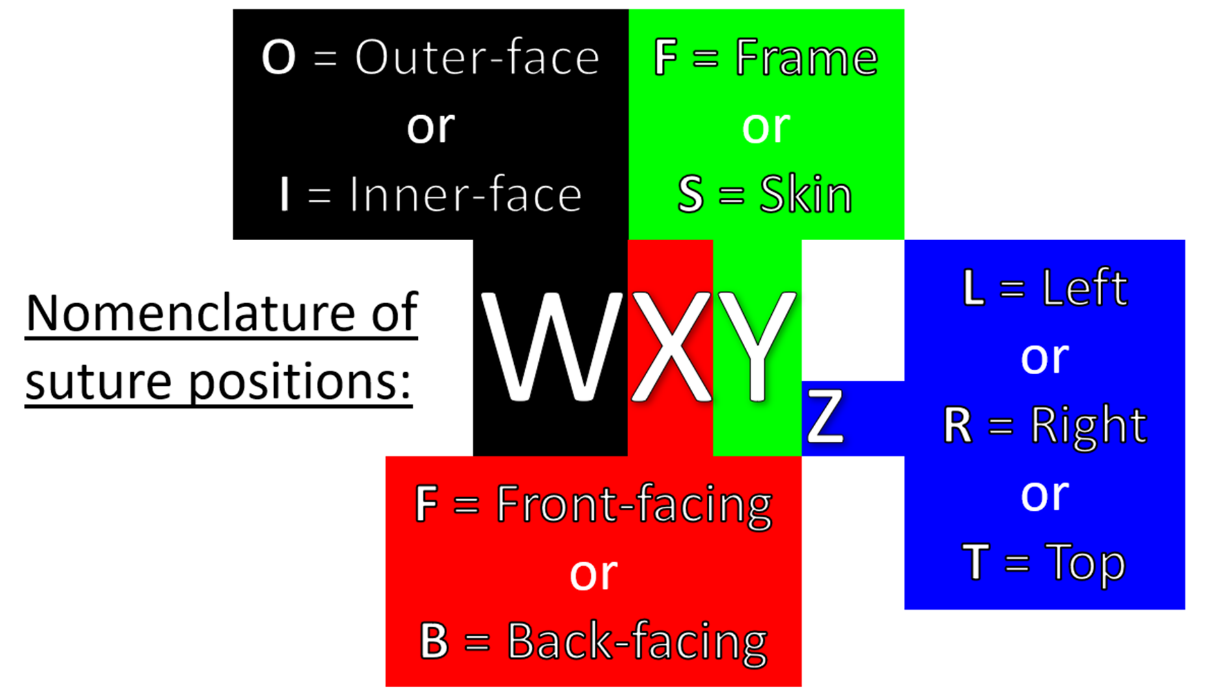

a. From outside the body (front side): OFS_T_ 🡪 IFS_T_ 🡪OFF_T_;
b. From inside the body: IFF_T_🡪 IBS_T_ (at this stage only the top of the front DSWC is partially underneath the skin as in **Figure** 2**k)**);
c. From outside the body (back side): OBS_T_ 🡪 IBF_T_ 🡪 OBF_T_ 🡪 OBF_L_🡪 IBF_L_🡪 OBS_L_;
d. From inside the body: IBS_L_ 🡪 IFS_L_ (at this stage the front frame should be approximately fully under the tissue, though without attempting to prop the chamber on top of the mouse’s back yet, as in **Figure** 2**l)**);
e. From outside the body (front side): OFS_L_ 🡪 IFF_L_ 🡪 OFF_L_ 🡪 OFF_R_ 🡪 IFF_R_ 🡪 OFS_R_;
f. From inside the body: IFS_R_ 🡪 IBS_R_;
g. From outside the body (back and front sides): OBS_R_ 🡪 IBF_R_ 🡪 OBF_R_ 🡪 Top of surgical guide;

Periodically flush the exposed dermis with warm saline to keep the wound hydrated.

Periodically apply tear gel to mice eyes.

Ensure mouse temperature within 24-30°C range.

20) Pull tautly both the leading end and the starting end of the suture from step 19)a. (OFS_T_), propping up the front and back frames and secure the frames together parallel to one another by tying the two suture ends.

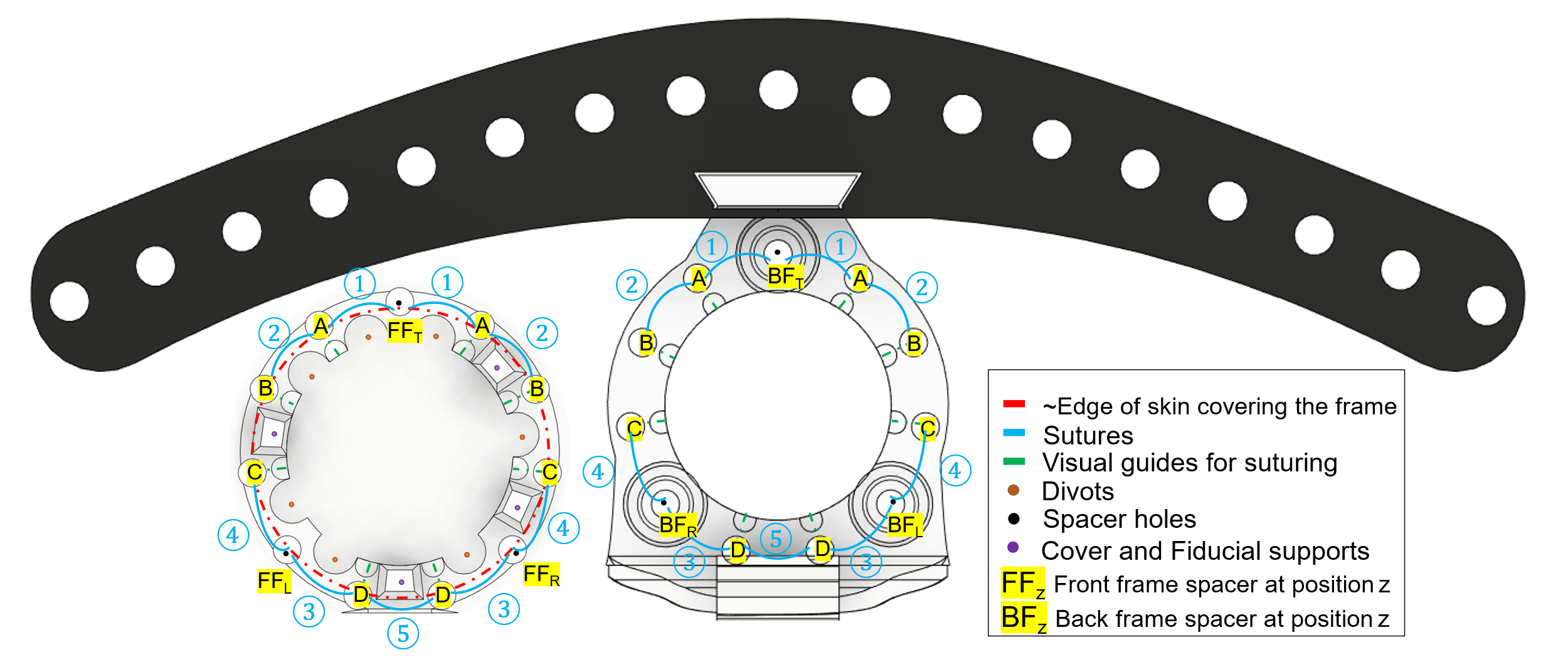

**Figure 3:**Annotated Front frame (left) and back frame in the surgical guide (on right) of DSWC.

21) Suture the frames together in pairs in the order numbered in **Figure 2**. Sutures ① through the top spacer followed by sutures ② through the neighbouring holes help ensure that the skin in the chamber is sufficiently tensioned with minimal gapping against the surface of the window, and that the skin on top of the frame is sufficiently cushioned to improve DSWC longevity. The remaining sutures help ensure proper seal of the DSWC with suture ⑤ being optional, As the suture holes on the front frame side should be underneath the skin, the frames have extrusions which may act as visual landmarks to find the holes (see **Figure 3**). These sutures should be performed sequentially through the covering skin, front frame, back skin and finally the back frame and back the opposite order through the neighbouring hole. These sutures knots should ideally be positioned on the front side frame in order for them to be protected by the chamber protector (added in **Figure** 2**t)**).
22) Carefully remove criss-cross temporary suture using scissors and forceps.
23) OPTIONAL: Cover the sutures in biocompatible resin and polymerize with UV light (while covering tissue with small piece of aluminum foil) or by applying Vetbond (biocompatible cyanoacrylate glue) to prevent mice from damaging sutures. N.B. Addition of these sealants prevent all future maintenance of the DSWC if deemed necessary.
24) Add hula clip support in back frame inserts and disconnect surgical guide. OPTIONAL: Apply Vetbond to hula clip to ensure it is properly secured. See **Figure 2u).**

Post-Procedure

1) Once surgery is complete, give mouse Metacam (5mg/kg, 0.5mg/mL) (subcutaneous injection). It is recommended to continue administration every ~24 hours for 48-72 hours to improve analgesia in this crucial recovery period (reducing mouse irritation translating into damaging of the DSWC).
2) Expose mouse to pure oxygen at 1 L/min and keep them warm while you wait for them to become ambulatory before returning to the cage.
3) Take note of time under anesthesia, dosages used and any notes about mouse recovery
4) Before returning the cage to the IVC racks.
   a. Fill out a SURGICAL CARD for your animal resources care facility.
   b. Print new cage card identifier as necessary.
