## Supplementary material for "Low-Cost 3D-Printed Tools Towards Robust Longitudinal Multi-Modal Pre-Clinical Imaging": Suppl 3 Surgical stage construction

To facilitate the delicate and high-precision dorsal skinfold window chamber installation procedure under isoflurane anesthetic, a novel surgical stage was designed and constructed.

Materials:

| 3D-printing:   - Fusion deposition modeling (FDM) 3D printer - High temperature- and chemical-resistance 3D printing filament (e.g., carbon-fiber – nylon (CF-Nylon) blend, or acrylonitrile styrene acrylate (ASA))   Adhesives:   - Acrylonitrile or polyurethane glue - PVA for Hot glue gun - Thread-locking fluid | Tools:   - Hot glue gun - Screwdrivers - Soldering iron - Wire-strippers   Apparatuses:   - 3D-printed surgical guide and surgical stage base - 5V 2A AC-DC power supply - Electric heating pad (nichrome wire) - Female DC connector (5.5mmx2.5mm) - Light-weight pen-light - Third-hand soldering stage Kotto^®^ |
| --- | --- |

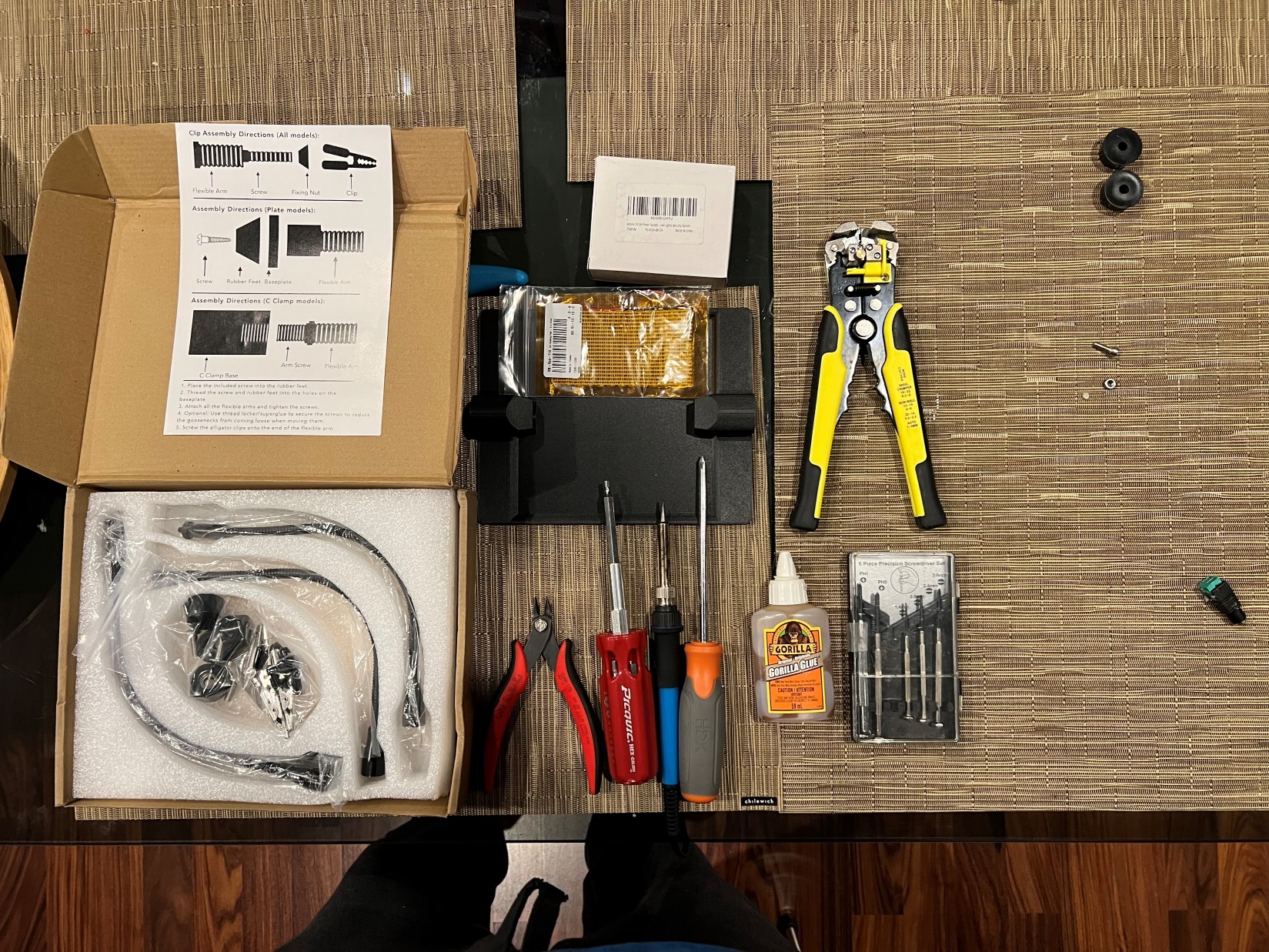

Figure 1: Materials required for assembly of surgical stage.

### Fabrication

Once the materials have been acquired and the surgical surface printed as seen in **Figure 1** & **Figure 2a)**, full assembly may begin (please refer to **Figure 2** for visual reference on steps):

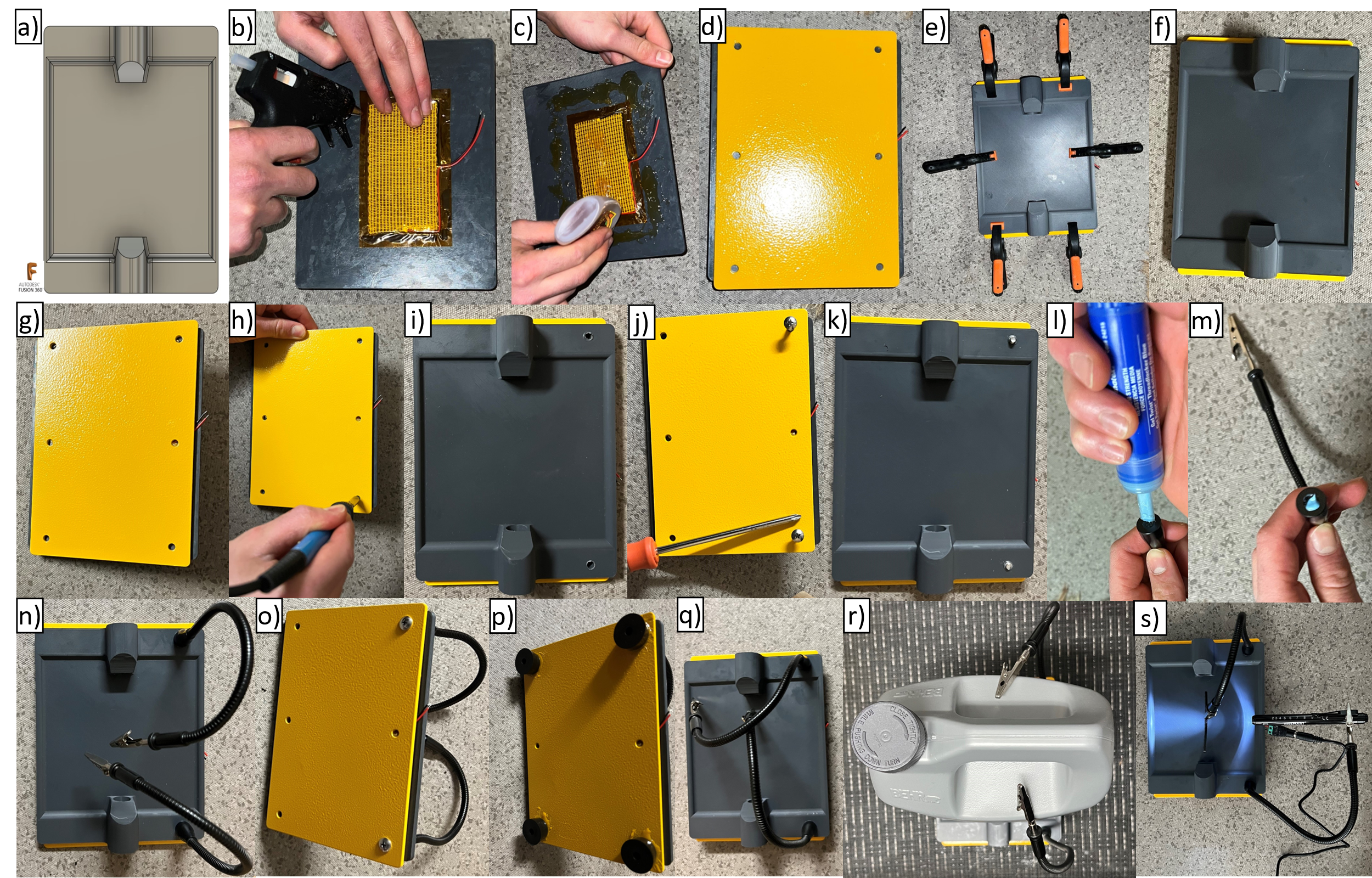

Figure 2: Outline of steps in assembly of surgical stage.

1. The surgical surface is flipped onto its backside to position and glue the electric heating pad just off-center, as in **Figure 2b)**. Hence the surgical surface may be half-heated, while the other half may remain relatively cool (thus during lengthier procedures where temperature regulation may be particularly challenging the mouse may be interchangeably transferred between cool and actively warmed surfaces).
2. Apply acrylonitrile or polyurethane glue to backside to adhere surgical surface to third-hand metal plate, as in **Figure 2c)-d)**.
3. Clamp surgical surface to third-hand metal plate, as in **Figure 2e)**. After leaving overnight, the surgical stage base should now be secured, as in **Figure 2f)-g)**.
4. Using the soldering iron, melt the surgical stage base through the hind bolt holes (where the electric heating pad wiring is protruding, as in **Figure 2h)-i)**.
5. Push the M4 bolts (which come packaged with the third-hand stage) through the backside of the surgical stage base, as in **Figure 2j)-k)**.
6. Apply thread-locking fluid to the inside of gooseneck alligator clip base-end, as in **Figure 2l)-m)**.
7. Tighten gooseneck alligator clip against surgical stage base, as in **Figure 2n)-o)**.
8. Adhere non-slip rubber feet to the backside of the surgical stage using acrylonitrile or polyurethane glue, as in **Figure 2p)-q)**.
9. Apply weight and leave to set overnight, as in **Figure 2r)**.
10. Secure female DC connector (5.5 mm x 2.5 mm) to electric heating pad wires, using wire-strippers and screwdriver as needed. At this point, the surgical guide, pen-light and 5V 2A AC-DC power supply may be connected completing the surgical stage, as in **Figure 2s)**.
