## Supplementary material for "Low-Cost 3D-Printed Tools Towards Robust Longitudinal Multi-Modal Pre-Clinical Imaging": Suppl 4 Novel OCT-histology corregistration pipeline

Suppl. 4: Novel OCT-Histology corroboration sacrifice and imaging protocol

Up to 2.5 months post-installation, depending on the window chamber and tumour condition, the mouse bearing the novel DSWC may be sacrificed for histology thus providing one of the gold-standard mechanistic corroboration for the intravital imaging findings. In the following document, the steps for co-registration between 2D histology and 3D imaging (e.g., Optical coherence tomography (OCT)) are outlined.

### 1 On the day of sacrifice (see steps visualized in **Figure 2**)

1. Mice are OCT imaged while anesthetized via isoflurane as per usual. If an exogenous staining agent is required for later histological analysis, please follow steps 2. - 3.
2. Mice are placed under heat lamp to ensure vasodilation (no anesthesia).
3. Mice will receive the exogenous agent via recommended route of administration. For example, intravenous agents may be administered via tail-vein injection using the custom-printed restrainer^1^ (see **Figure 1**).

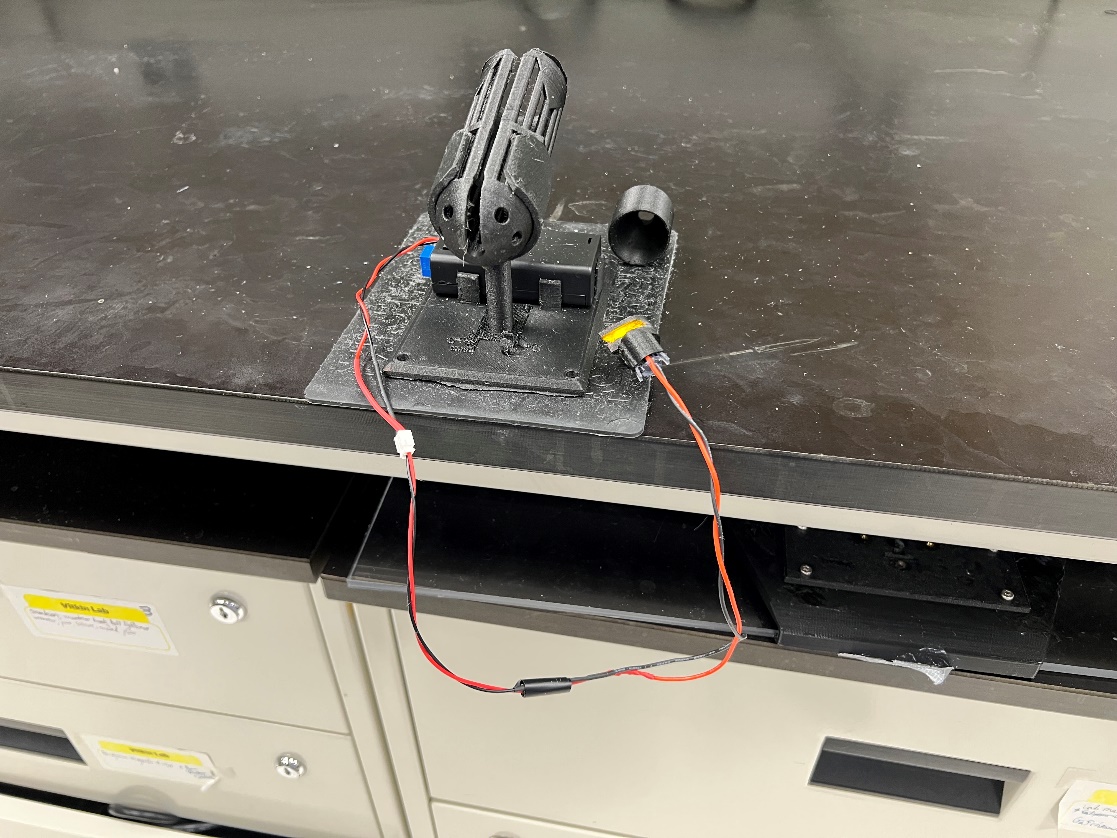

**Figure 1: Tail-vein injection setup with yellow LED light guiding more rapid and accurate tail-vein injections, important for such time and dose-sensitive studies.**

In the special case where this is not possible (due to damage to the tail-vein), retro-orbital injections will be performed after administration of Alcaine ophtalmic local anesthesia.

4. ~15min prior to sacrifice, mice will be anesthetized via ketamine/xylazine cocktail at concentrations (100mg/kg, 10mg/kg).
5. Mice are sacrificed via cervical dislocation.
6. The full window chamber is excised and quickly placed into 10% formalin for ~48 hours.

### 2 48 hours post-sacrifice (see steps visualized in **Figure 2**)

7. After ~48 hours have elapsed, remove the tissue and allow to air-dry (avoid having to transfer the DSWC to 70% isopropyl alcohol as this results in its distortion).
8. OCT image the excised DSWC frame, thus obtaining a structural scan of the tissue prior (step 1.) and post fixation, allowing tracking of the various tissue distortions^2^.
9. Insert the histology fiducials tool (**Figure 2**(c)-(d)) on the front-frame of the OCT.
10. Redo the OCT imaging with the fiducials tool for lateral co-registration to OCT to be complete.
11. Carefully remove the sutures retaining the front-frame of the DSWC before removing the front-frame, taking care to keep track of the center of the impressions left by the spacers. These can be marked with surgical marker if deemed necessary.
12. Remove the back DSWC frame.
13. Tightly Suture the histology stencil tool (**Figure 2** (g)-(h)) such that the tissue is under sufficient tension to enable proper marking without tearing the fixed tissue.
14. Mark “VI pattern” onto the side of the exposed panniculus carnosus muscle^3–5^ using 2 colours of tissue marking dye in order to later distinguish transverse orientation of tissue sections along OCT’s fast-scanning x-axis (<https://www.cancerdiagnostics.com/consumables/grossing/tissue-marking-dye>).
15. Slice the tissue parallel to the plane of B-scan acquisition, according to the horizontal slots (**Figure 2** (g)), with a third dye mark the bottom of the middle tissue section to prevent confusion on transverse orientation along OCT’s slow-scanning y-axis.
16. Paraffin embed the sectioned tissue and prepare for immunohistochemistry staining.

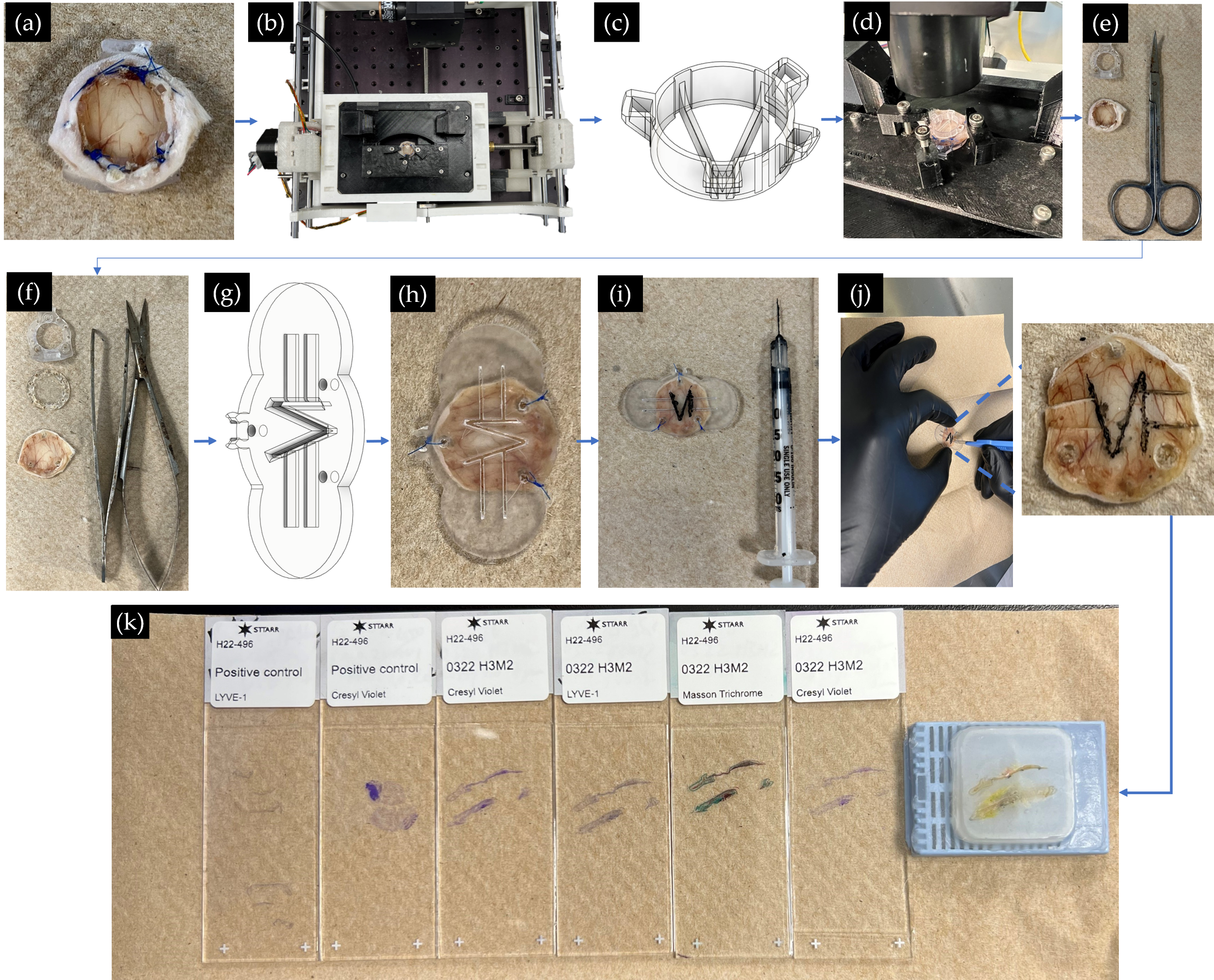

**Figure 2: Novel OCT-histology co-registration pipeline.** (a) Excised DSWC after 48-72h of 10% formalin fixation (but no more, to reduce masking of epitopes and cross-linking of antigens). This chamber is then OCT imaged structurally (b) pre and (d) post placement of (c) histology fiducials tool. (e) & (f) The DSWC sutures and frames are then removed and the histology stencil tool pictured in (g) is sutured through the previous suture holes of the spacers as in (h). (i) Tissue marking dye is traced via 29G need through the stencil. (j) A scalpel is used to perform preliminary tissue sections parallel to OCT B-scans (a zoomed in-view of the tissue following removal of histology stencil tool is shown). (k) Paraffin-embedded tissue sample and histology tissue sections with visible dots for guiding OCT-histology co-registration.

At this stage, histology sections may be co-registered to OCT (or potentially other high resolution modality, following the data acquisition protocol described above), using the custom algorithms scripted in MATLAB, which can be found in ^6^.

We note here that previously we were interested in distinguishing chronic hypoxia from acute hypoxia through the administration of a marker of global hypoxia pimonidazole HCl, and of a perfusion agent, known as Hoechst 33342 (administered by tail vein injection with 100uL at 10mg/mL in 0.9% saline 1min prior to sacrifice)^7^. However due to the unlikelihood of observing a change in vessel diameter along the lines of acute hypoxia tumour vessel shunting, this will not be performed. Although it is possible that via automation of OCT angiography post-processing by implementing algorithms such as ID-BISIM^8^, we may potentially achieve sufficient accuracy to distinguish this vessel diameter change from noise, given the resolution limits of our OCT system, the logistical complications of performing repeated OCT imaging to be able to capture the changes in perfusion over time prior to sacrifice may jeopardize the survival of the specimen before the foreseen time of death, which is a too large risk given the importance of this final timepoint.
