## Supplementary material for "Low-Cost 3D-Printed Tools Towards Robust Longitudinal Multi-Modal Pre-Clinical Imaging": Suppl 5 DSWC and toolset animation

Suppl. 5: Video file showcasing novel dorsal skinfold window chamber and peripheral imaging toolset designs

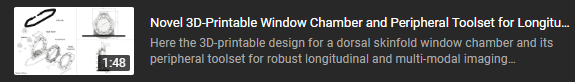

<https://youtu.be/9fJpV3muag0>
