## Supplementary material for "Low-Cost 3D-Printed Tools Towards Robust Longitudinal Multi-Modal Pre-Clinical Imaging": Suppl 6 Automatic xy-microstage in practice

Suppl. 6: Video file representing 3D printed automatic xy-microstage in practice

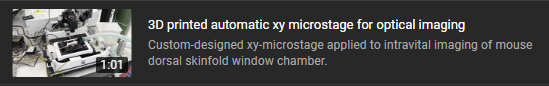

<https://youtu.be/51pJqcYvFUQ>
